# Membrane and glycocalyx tethering of DNA nanostructures for enhanced uptake

**DOI:** 10.1101/2023.03.09.529286

**Authors:** Weitao Wang, Bhavya Chopra, Vismaya Walawalkar, Zijuan Liang, Rebekah Adams, Markus Deserno, Xi Ren, Rebecca E. Taylor

**Author notes:** indicates equal author contribution (Weitao Wang, Bhavya Chopra, Vismaya Walawalkar, Rebecca E. Taylor).

## Abstract

DNA nanostructures (DNs) have been increasingly utilized in biosensing, drug delivery, diagnostics and therapeutics, because of their programmable assembly, control over size and shape, and ease of functionalization. However, the low cellular uptake of DNs has limited their effectiveness in these biomedical applications. Here we demonstrate the potential of membrane and glycocalyx binding as general strategies to enhance the cellular uptake of DNs. By targeting the plasma membrane and cell-surface glycocalyx, the uptake of all three distinct DNs is significantly enhanced as compared to uptake of bare DNs. We also demonstrate the viability of single-step membrane labeling by cholesterol-DNs as competitive with previous multistep approaches. Further, we show that the endocytic pathway of membrane-bound DNs is an interdependent process that involves scavenger receptors, clathrin-, and caveolin-mediated endocytosis. Our findings may potentially expand the toolbox for effective cellular delivery of DNA nanostructured systems.

## Introduction

Nanoscaled platforms have drawn enormous research effort in the past decade for imaging, biosensing and drug delivery.^1–8^ These nanoplatforms can carry molecular loads, including diagnostic and therapeutic agents, serving as effective delivery vehicles to target specific cell-types and intracellular compartments.^9,10^ Nevertheless, their internalization and accumulation inside of cells have raised concerns over cytotoxicity.^11–13^ Moreover, the need for personalized therapeutics and diagnosis requires versatile and programmable delivery systems, which limits our choice of delivery vehicles.^14,15^

Structural DNA nanotechnology is recognized as a practicable candidate to potentially overcome the aforementioned obstacles because of its biocompatibility, biodegradability, programmability and ease of functionalization.^7,8,16–21^ Since its emergence, various DNA nanostructures (DNs) have been designed and synthesized with nanoscale precision, and modified with chemical and biological conjugates.^20,22–25^ For example, recent studies show the potential of DNs to enable robust expression genetic cargo for therapeutics.^8,26–30^ Despite the myriad of achievements made, one challenge that remains is the low cellular uptake of DNs.^31^ Previously, the cellular uptake of DNs has been studied in terms of geometric parameters like size and shape, wireframe architecture, molecular weight, and surface chemistry.^21,32–37^ For example, Bastings *et al*. reported that compact and solid DNs with low aspect ratios resulted in substantially higher DN uptake.^32^ Moreover, DNs could trigger cellular uptake via interactions with specific membrane receptors.^31^ This has sparked further studies, such as programming DNs to bind to membrane receptors to activate endocytosis, and decorating DNs with ligands, such as aptamers, peptides and proteins to improve DN uptake.^38–42^ However, taking these factors into consideration, the concentration of internalized DNs may still be insufficient to fulfill their therapeutic capabilities. Therefore, there is a need develop strategies for enhancing the cellular uptake DN-based delivery systems.

Applying theoretical analysis and molecular dynamics simulations, previous studies have demonstrated that reducing the separation between substances and cellular membranes triggered stronger membrane-substance interaction, which facilitated the internalization.^43–45^ A more recent experimental study has suggested that a stronger membrane-DN interaction may lead to preferable DN uptake.^33^ These results raise the question of whether DN uptake can be enhanced by simply bringing DNs and cellular membranes into close proximity without harnessing the additional peptides or ligands. Limited studies appear to support such a hypothesis. For example, 6-duplex nanobundles modified with cholesterol anchors had significant enhancement in their uptake as compared to unmodified nanobundles.^46^ Binding DNA nanotiles onto the cell-surface glycocalyx also enhanced nanotile’s uptake.^47^ While these studies provide new insights into the relation between an enhanced membrane-DN interaction and an increase in NP uptake, the topic has not been fully explored, and many important questions need to be answered, for instance, if the relation between a stronger membrane-DN interaction and a higher uptake applies to a broader range of DNs, and how to engineer and optimize such an enhanced interaction. Further, it is unclear if different methods of immobilization will be subject to different endocytic pathways.

Toward this end, here we performed a comprehensive study with the aim of understanding the relation between the membrane and glycocalyx binding of DNs and the their uptake. Cell surface-bound DNs and internalized DNs were both qualitatively visualized and quantitatively measured using previously reported methods.^47,48^ We first revisited how uptake was affected by DN geometry using DNA nanospheres, nanorods and nanotiles.^47,49,50^ By modifying DNs with membrane binding moieties, cholesterol-mediated membrane anchoring was performed using both a two-step and recently developed one-step method while glycocalyx anchoring was performed through hybridization to click-anchored ssDNA. We find for all case that cell surface anchoring significantly enhanced the uptake of DNs as compared to unmodified DNs (up to 8-fold). We conducted further research into the impact of binding moiety valency and placement on DN uptake and found that both factors played a significant role in enhancing uptake. These studies also establish the utility of directly conjugating cholesterol onto DNs for applications in which multistep methods are impractical. Finally, we explored the feasibility of membrane wrapped-mediated uptake and evaluated endocytic pathways of membrane-bound DNs. These findings provide new insights into the membrane tethering-enhanced cellular uptake. Our approach may strengthen and expand the toolkit for leveraging cellular uptake in DN-delivery based therapeutic applications.

## Results and discussion

### Design, Synthesis and Characterization of DNs

Three shapes of DNs, including DNA nanospheres (S, *∼*54 nm in diameter), nanorods (R, *∼*7 nm in diameter and *∼*400 nm in length) and nanotiles (T, 96 nm x 74 nm), were designed and synthesized (Fig. 1A).^49,50^ All DNs were labeled with 14 biotin molecules for subsequent fluorescence staining. In order to investigate the effect of valency and placement of single-stranded DNA (ssDNA) overhang on cellular uptake, each shape of DNs was modified with three types of ssDNA overhang decorations: 0 overhangs (S0, R0, T0), 10 overhangs uniformly presented on the central surface (S10, R10, T10), and 4 overhangs with each edge of the DN having 2 overhangs (S4, R4E, T4E. Since a sphere doesn’t differ center and edge on its surface, we used S4 instead S4E). All DNs were characterized with atomic force microscopy (AFM) and agarose gel electrophoresis, which confirmed their formation.

**Figure 1:**
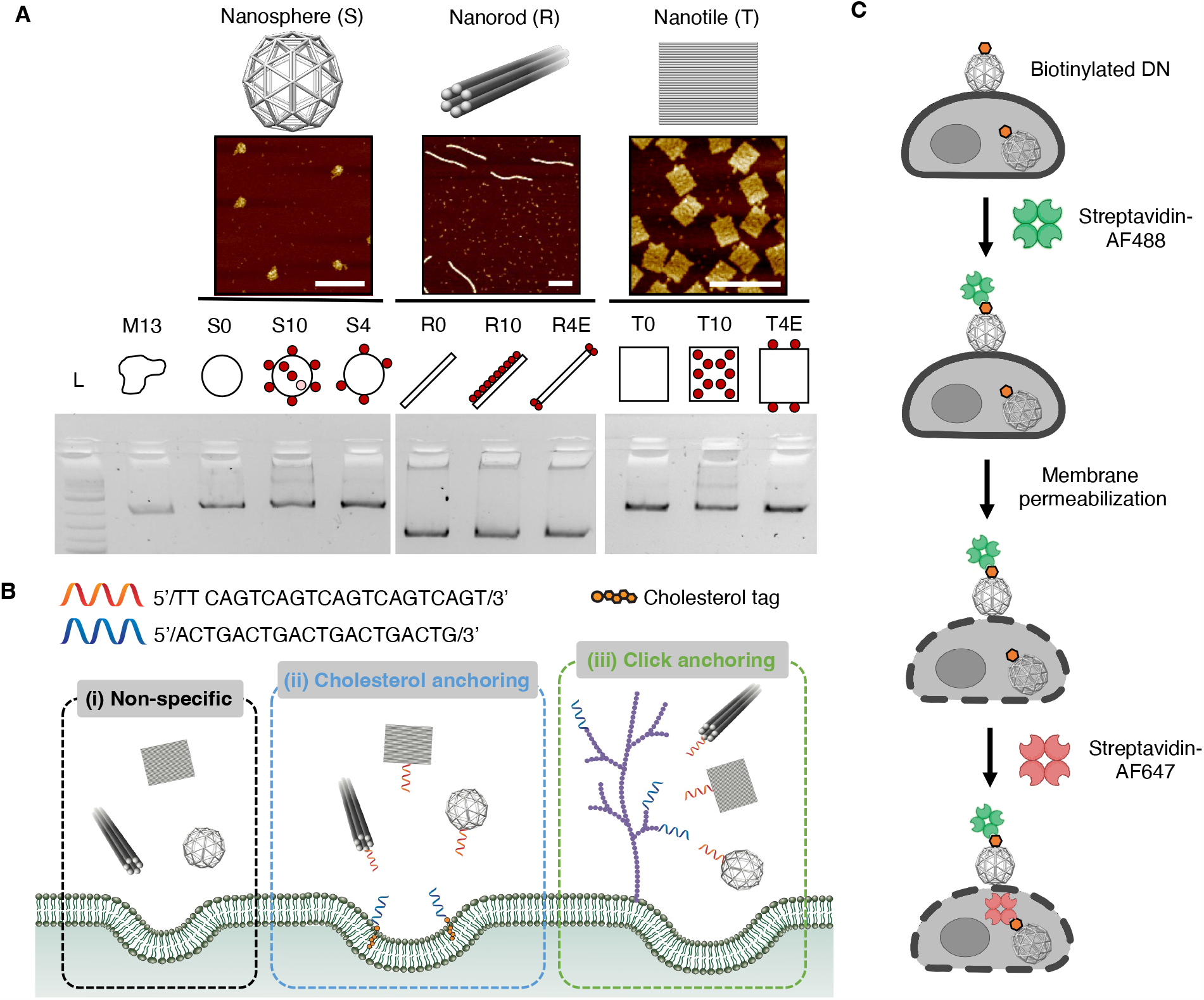
Design and characterization of DNs, the membrane binding pathways and dual-color streptavidin staining of membrane-bound and internalized DNs. (A) Schematic illustrations, AFM characterizations and agarose gel electrophoresis of DNA nanospheres (S), nanorods (R) and nanotiles (T). All scale bars: 200 nm. (B) Three membrane binding strategies, including (i) non-specific adhesion of DNs with no overhangs; (ii) two-step cholesterol membrane anchoring though the hybridization of DN-surface ssDNA overhangs and membrane-immobilized cholesterol-ssDNA anchors; (iii) two-step click glycocalyx anchoring though the hybridization of DN-surface ssDNA overhangs and cell-surface glycocalyx-immobilized ssDNA anchors. (C) Dual-color streptavidin staining technique. After cell fixation, membrane-bound DNs were first stained with streptavidin-Alexa Fluor 488. Cellular plasma membranes were then permeabilized using Triton X-100. Lastly, streptavidin-Alexa Fluor 647 was administrated and stained the internalized DNs

### Membrane Binding-enhanced Uptake of DNs

Three strategies were studied to bind DNs on cell membranes (Fig. 1B). Bare DNs with no overhangs can only non-specifically adhere to membranes through a combination of interactions like electrostatic interactions. This condition was considered as a negative control. For overhang-decorated DNs, a two-step cholesterol membrane anchoring and two-step click glycocalyx anchoring were used.^47,51^ In two-step cholesterol anchoring, cell membranes were first immobilized with cholesterol-ssDNA anchors by spontaneously inserting the hydrophobic portion of the amphiphile into the lipid membranes. In a second step, DNs modified with the complementary ssDNA hybridized with the membrane-immobilized ssDNA anchors, leading to the recruitment of DNs onto the membrane. For click glycocalyx anchoring, ssDNA anchors were incorporated onto cell-surface glycocalyx through metabolic glycan labeling and glycocalyx labeling with copper-free click chemistry. Then DNs carrying complementary ssDNA hybridized with glycocalyx-immobilized ssDNA anchors, leading DNs to attach onto the glycocalyx.

Second is the staining technique. In order to visualize both the membrane-bound DNs and internalized DNs, we utilized a dual-color streptavidin staining method (Fig. 1C).^47^ Cells were first fixed after incubating with biotinylated DNs. Next, streptavidin-Alexa Fluor 488 (AF488) was introduced to stain the membrane-bound DNs through the conjugation of biotin and streptavidin, followed by the permeabilization of cellular plasma membrane. Then internalized DNs were stained by administering streptavidin-Alexa Fluor 647 (AF647). Fluorescence microscopy images were taken to assess the membrane binding and uptake of DNs. For quantitative measurement, in order to lower the impact from the noise caused by non-specific internalization of fluorophores, we applied our previously reported computational pipeline to remove undesired background and capture the signals of internalized DNs more precisely (Fig. S1).

To answer the question of whether membrane binding enhances the cellular uptake of DNs, we administered 9 DNs into an adherent mammalian cell line, human umbilical vein endothelial cells (HUVECs), separately, for 30 min incubation at 37°C. The cell-surface and internalization signals were measured, and an internalization efficiency was defined by dividing internalization signal intensity by the sum of internalization and on-membrane signal intensities (Eq. 1).

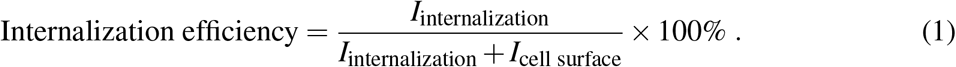

The efficiency was defined in this way to account for the influence of membrane-bound DNs. We first revisited the geometry effect by comparing the non-specific membrane adhesion and uptake of DNs with no overhangs. We found nanospheres outperformed nanorods and nanotiles in both cell-surface and internalization signal intensities, which agreed with previous studies (Fig. 2A and 2B).^32,33^ Surprisingly, though having low signal intensities, nanotiles possessed a higher internalization efficiency as compared to two other shapes. To study the effect of overhangs on membrane binding and internalization, the readouts of DNs with overhangs were normalized to bare DNs in each shape. For instance, S10 and S4 were normalized to S0 in cell-surface, internalization and efficiency plots. All three shapes of DNs with overhangs experienced significant enhancement in their membrane binding and internalization using cholesterol anchoring (Fig. 2C and 2D). For example, nanospheres S10 had a 5-fold increase in cell-surface signal intensity and a 2 to 3-fold increase in internalization signal. The placement of overhangs appeared to greatly impact not only membrane binding of the DNs, but also their uptake. In particular, edge-decorated nanotiles T4E had a 4-fold increase in both cell-surface and internalization signal intensities as compared to center-decorated nanotile T10, and it was more preferentially internalized than S10 (Fig. S2). Surprisingly, however, with the enhancement in the uptake of DNs with overhangs, their internalization efficiency dropped compared to DNs with no overhangs. The trend held for all shapes, presumably because the cellular uptake efficiency had plateaued while DNs in the buffer continued to bind to membranes. These findings support our assumption that using cholesterol anchoring technique to bind DNs onto cell plasma membranes, their cellular uptake can be enhanced.

**Figure 2:**
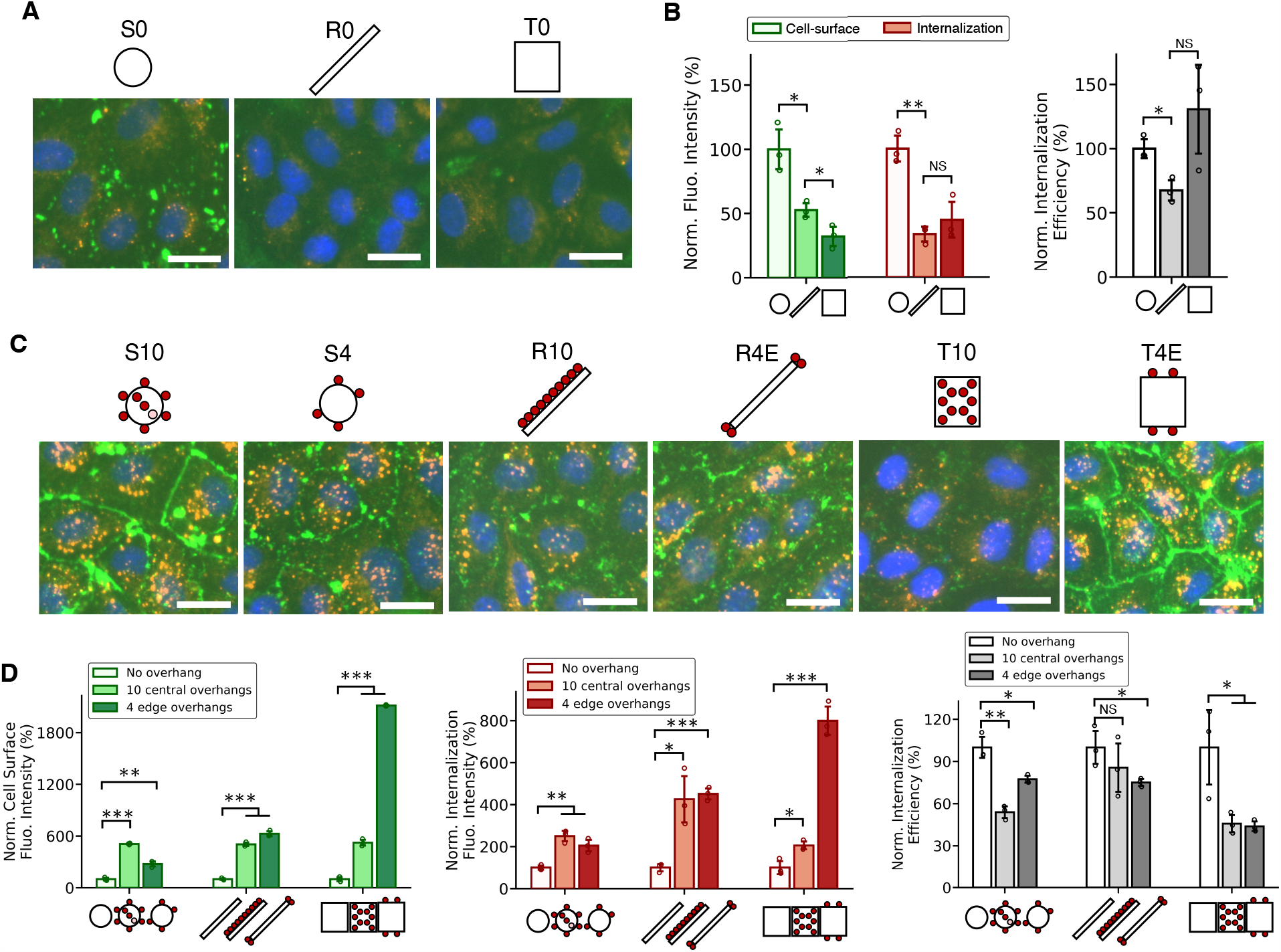
Cellular uptake analysis of DNs with membrane binding ability via two-step cholesterol membrane anchoring. (A) Fluorescence microscopy images of HUVECs after incubating with DNs with no overhangs for 30 min at 37°C. (B) Quantification of fluorescence intensities of cell-surface signal, internalization signal and internalization efficiency of DNs with no overhangs. In all three categories, the data were normalized to the readouts of nanospheres. (C) Fluorescence microscopy images of HUVECs after incubating with DNs with overhangs for 30 min at 37°C. (D) Quantification of the fluorescence intensities of cell-surface signal, internalization signal and internalization efficiency of DNs with overhangs. In each shape, the data were normalized to DNs with no overhangs. Specifically, S10 and S4, R10 and R4E, T10 and T4E were normalized to S0, R0 and T0, respectively. For all fluorescence microscopy images, membrane-bound DNs were stained with streptavidin-AF488 (green) and internalized DNs were stained with streptavidin-AF647 (red). Cell nuclei were stained with Hoechst (blue). Data were presented as means ± s.d. in (B) and (D) with n=3. \**P ≤* 0.05, \*\**P ≤* 0.01, \*\*\**P ≤* 0.001. All scale bars: 10 µm

To investigate if other membrane binding strategies also lead to an enhancement in DN uptake, we turned to click glycocalyx anchoring in which DNs were anchored onto cell-surface glycocalyx. In this case, the non-specific membrane binding and internalization of DNs with no overhangs seemed to be lower than they were in the control groups of cholesterol anchoring, potentially due to the modification of glycocalyx (Fig. 3A and 3B). Both cell-surface and internalization signals with overhangs increased similarly in click anchoring as in cholesterol anchoring (Fig. 3C and 3D). The valency and position of overhangs were shown to be important, again, by comparing nanorod R4E and R10, and nanotile T4E and T10. Here T4E had the most internalization among all structures as well (Fig. S3). To briefly summarize, we observed a consistent relation between a significant enhancement in membrane-bound DNs and a similar increase in internalized DNs, in both cholesterol membrane anchoring and click glycocalyx anchoring, which shows the general appliance of the strategy that utilizes cell surface binding for uptake enhancement.

**Figure 3:**
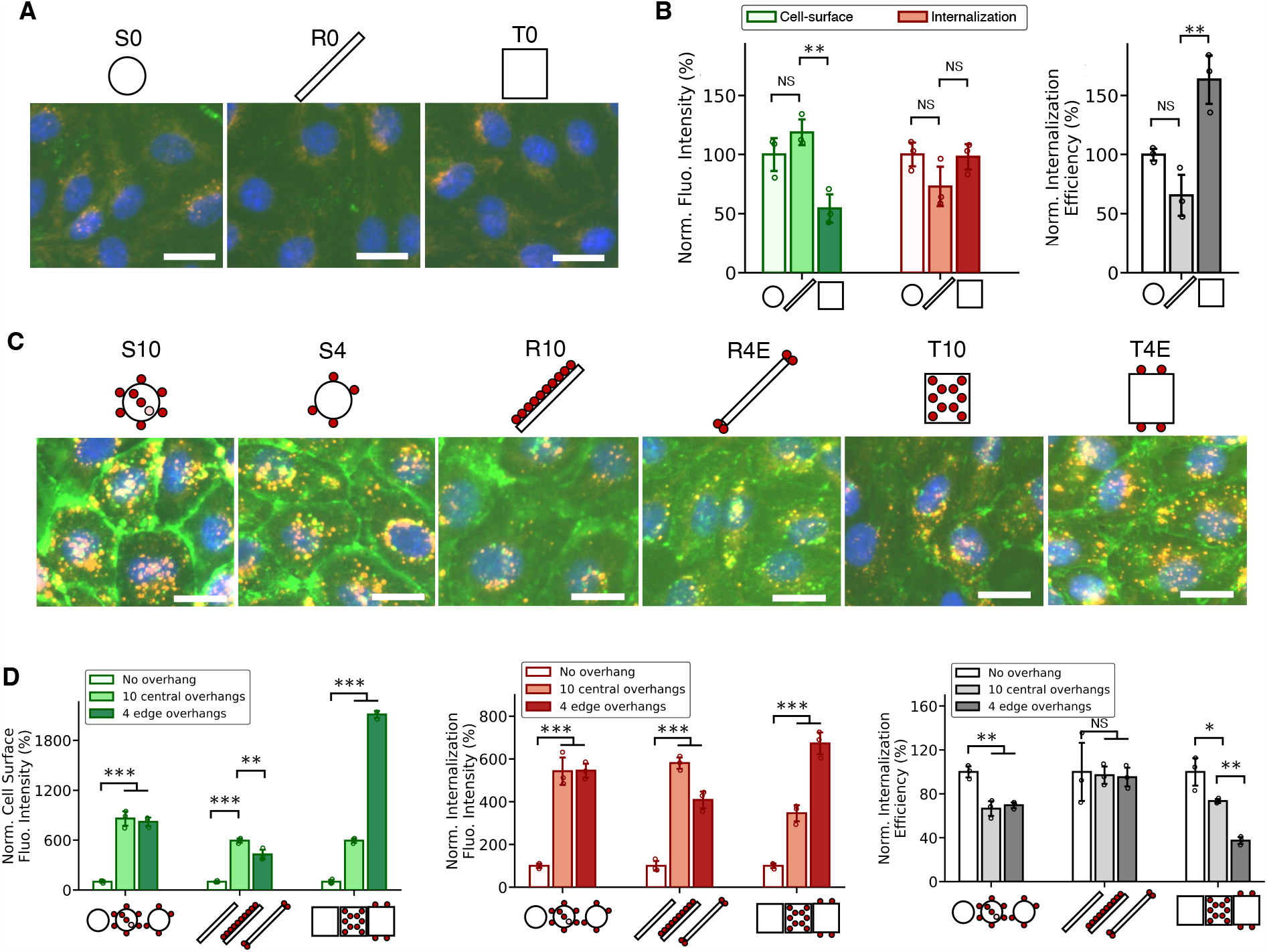
Cellular uptake analysis of DNs with membrane binding ability via two-step click gly-cocalyx anchoring. (A) Fluorescence microscopy images of HUVECs after incubating with DNs with no overhangs for 30 min at 37°C. (B) Quantification of fluorescence intensities of cell-surface signal, internalization signal and internalization efficiency of DNs with no overhangs. In all three categories, the data were normalized to the readouts of nanospheres. (C) Fluorescence microscopy images of HUVECs after incubating with DNs with overhangs for 30 min at 37°C. (D) Quantification of the fluorescence intensities of cell-surface signal, internalization signal and internalization efficiency of DNs with overhangs. In each shape, the data were normalized to DNs with no over-hangs. Specifically, S10 and S4, R10 and R4E, T10 and T4E were normalized to S0, R0 and T0, respectively. For all fluorescence microscopy images, membrane-bound DNs were stained with streptavidin-AF488 (green) and internalized DNs were stained with streptavidin-AF647 (red). Cell nuclei were stained with Hoechst (blue). Data were presented as means ± s.d. in (B) and (D) with n=3. \**P ≤* 0.05, \*\**P ≤* 0.01, \*\*\**P ≤* 0.001. All scale bars: 10 µm

We further investigated other factors that might affect the uptake of DNs, including incubation time, the concentration of DNs and cholesterol anchors, and the buffer for incubation, using three edge-decorated DNs, S4, R4E and T4E. We noticed that while membrane-bound DNs continued to increase with longer incubation time, the internalization signal appeared to plateau with no significant changes for all three shapes (Fig. S4). However, if the cells were first incubated with DNs, then washed and followed by another incubation, we saw a rapid decrease in cell-surface and internalization signals. We postulated that after washing, there were no DNs in the buffer available to continue binding to cell membranes, thus limiting the membrane binding internalization of DNs.

The concentration of DNs and cholesterol anchors positively affected the membrane binding and uptake of DNs, as previously reported.^32,35^ By changing DN concentration from 1 nM, 5 nM to 10 nM, or the concentration of cholesterol anchors administrated to cells, from 0.1 µM, 0.5 µM to 1 µM, we observed a monotonic increase in both the on-membrane and internalization signal intensities with an increasing concentration of DNs or cholesterol anchors (Fig. S5 and Fig. S6). However, it is worth noting that too high of a concentration of cholesterol anchors resulted in substantial aggregation of DNs on cell membranes (Fig. S6).

DNA nanostructures, especially DNA origami, rely on positive ions in the buffer to maintain their structures.^16–18,21^ This leads to the question of whether the buffer has an influence on the membrane binding and uptake of DNs. Here we used four buffers, including phosphate buffered saline (PBS), roswell park memorial institute (RPMI) 1640 medium, endothelial cell growth basal medium-2 (EBM-2) and endothelial cell growth medium-2 (EGM-2). We found DNs in PBS buffer were most preferably bound to cell membranes and internalized than other cell medium (Fig. S7). DNs in EGM-2 were the least bound to cell membranes and internalized. We reasoned fetal bovine serum (FBS) in EGM-2 might be the cause, which destabilizes DNs.

### One-step Cholesterol Anchoring versus Two-step Cholesterol Anchoring

Lipophilic cholesterol tags have been widely used as a modification to DNs, enabling DNs to interact with and bind to cellular membranes.^52–55^ Yet the modification with hydrophobic cholesterol tags on DNs may lead to undesired aggregation of DNs, damaging their structural integrity and thus limiting the applications.^56,57^ It is for this reason, the two-step cholesterol anchoring method was developed to avoid the aggregation problem.^22,47,51^ Recently, Ohmann et al. have demonstrated that a strategic choice of cholesterol-conjugated sequences can successfully control the aggregation of cholesterol-modified DNs.^56,57^ To place cholesterol TEG linkers on an overhang sequence, incorporating an ssDNA overhang adjacent to the cholesterol and thus presenting an shielding effect is able to control the aggregation and increase the yield of one-pot cholesterol-conjugated DNs. Studies on lipid vesicles, such as GUVs or SUVs, have shown that the spontaneous insertion of cholesterol into lipid membranes was not impeded by the presence of a shielding overhang in the one-pot assembly of cholesterol-conjugated DNs. The successful synthesis of cholesterol-conjugated DNs opens new possibilities for *in vivo* research. Therefore, we think it is important to investigate the difference in the membrane binding and cellular uptake between DNs with direct conjugation of cholesterol tags, which we refer as the one-step cholesterol anchoring, and the two-step cholesterol anchoring, to provide guidance in the future applications of cholesterol-modified DNs (Fig. 4A).

**Figure 4:**
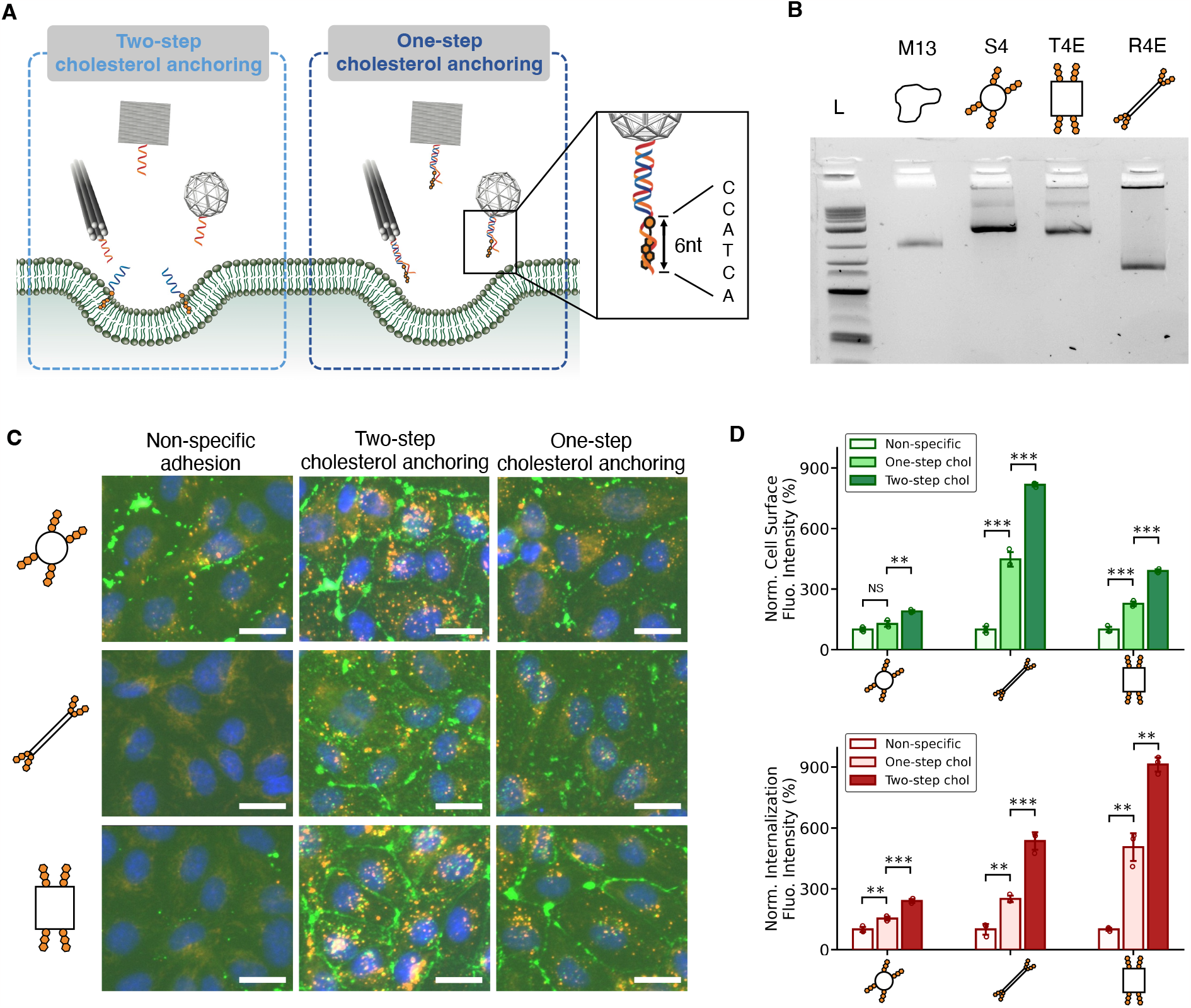
One-step cholesterol anchoring and two-step cholesterol anchoring on the membrane binding and cellular uptake of DNs. (A) Schematic illustrations of one-step and two-step cholesterol anchoring. Cholesterol molecules were directly conjugated onto DNs with a shielding ss-DNA overhang adjacent to the cholesterol. (B) Agarose gel electrophoresis of DNA nanospheres, nanorods and nanotiles conjugated with four cholesterol molecules on their edges. (C) Fluorescence microscopy images of HUVECs after incubating with DNs for 30 min using one-step and two-step cholesterol anchoring. Membrane-bound DNs were stained with streptavidin-AF488 (green) and internalized DNs were stained with streptavidin-AF647 (red). Cell nuclei were stained with Hoechst (blue). Scale bars: 10 µm. (D) Quantification of fluorescence intensities of cell-surface and internalization signals for DNs using one-step and two-step cholesterol anchoring. The data were normalized to bare DNs. Data were presented as means ± s.d. with n=3. \*\**P ≤* 0.01, \*\*\**P ≤* 0.001.

For this one-step approach, we modified nanosphere S4, nanorod R4E and nanotile T4E by directly conjugating cholesterol tags onto the edges and incorporating an additional complementary ssDNA for a shielding effect (Fig. 4A). The complementary ssDNA is 6 base-pairs longer than the cholesterol-conjugated sequence. Monodispersed bands were observed for all DNs in agarose gel electrophoresis with no significant aggregations, confirming the successful formation of DNs (Fig. 4B). Here again, fluorescence microscopy images of HUVECs showed preferential membrane binding and uptake of DNs with one-step cholesterol anchoring comparing to bare DNs (Fig. 4C). Nanospheres outperformed nanotiles and nanorods, demonstrating the geometry effect still held for the one-step approach. Intriguingly, compared to two-step cholesterol anchoring, the cell-surface and internalization signal intensities of one-step cholesterol anchoring were weaker. Considering that DNs in both methods had the same number of tags or overhangs, this finding suggests that the efficiency of spontaneous insertion of cholesterol into lipid membranes might be lower when the cholesterol is directly conjugated on DNs. Nonetheless, the highlevel of conjugation and uptake validate the one-step approach as a simple approach that is compatible with future approaches, such as *in vivo* studies, that may not accommodate washing steps.

### Endocytic pathways of membrane-bound DNs

The enhanced cellular uptake of membrane-bound DNs raised the question whether membrane attachment might facilitate the internalization of DNs via non-endocytic processes. To investigate whether this is plausible, we analyzed adhesion-driven particle envelopment using a simple elastic continuum description. Specifically, by modeling lipid membranes as fluid curvature-elastic surfaces, we calculated the energy needed for a membrane to wrap, engulf, and finally internalize a DN, and compared this to the energy available due to the insertion of cholesterol tags into the membrane (Fig. S8). We found that this adhesion energy was substantially smaller than what would be needed to drive passive uptake (Supplementary Table. 1). Take nanospheres as an example, the membrane bending energy to engulf a nanosphere is more than about 600*k*_B_*T*, thus it takes approximately 50 cholesterol tags for the membrane to spontaneously warp the nanosphere (please refer to Supplementary Information for more details on calculations). The number is way larger than what we have on our nanospheres, suggesting that DNs are internalized through active endocytosis process.

Using nanospheres with 4 overhangs (S4), we then sought to understand whether membranebound DNs had similar endocytic pathways as bare DNs, or whether there were new pathways involved. Scavenger receptors, clathrin-mediated endocytosis and caveolin-mediated endocytosis were reported to be responsible for internalizing various shaped DNs.^33,35^ Polyinosinic acid (Poly-I) is often used to bind and saturate scavenger receptors. First, cells were introduced with Poly-I at 400 µg*/*ml and incubated at 37°C for 30 min. For all three membrane binding strategies, including non-specific adhesion, two-step cholesterol anchoring and click anchoring, the uptake of S4 dropped 40%-50% as compared to nontreated cells, which demonstrated the importance of scavenger receptors in internalizing membrane-bound S4 (Fig. 5A and 5B). Interestingly, the cellsurface signals also reduced, showing the treatment of Poly-I affected the ability of S4 to bind to cell membranes. Next, the role of clathrin-mediated endocytosis was investigated by using Pitstop-2 and Dynasore as inhibitors. Pitstop-2 selectively blocks the recruitment of clathrin proteins to the plasma membrane, thereby preventing the formation of clathrin-coated pits and subsequent vesicle formation. We found cells treated with 10 µM Pitstop-2 resulted in only 10%-20% of internalized signals as compared to nontreated cells, suggesting the strong relation between clathrin and the uptake of membrane-bound S4 (Fig. 5C and 5D). Dynasore blocks the activity of dynamin so that the pinching off of endocytic vesicles from the plasma membrane is inhibited. A 50% decrease in S4 uptake was observed in cells treated with 120 µM dynasore (Fig. S9). Further, methyl-*β* -cyclodextrin (M*β* CD) is an inhibitor to caveolin-mediated endocytosis by depleting membrane cholesterol, disrupting the structure and function of caveolae. Its relation to S4 uptake appeared to be weaker, with only 10% and 20% decrease in non-specific adhesion and click anchoring, respectively (Fig. S10). However, the uptake of S4 in cholesterol anchoring had more than 40% decrease, presumably because of the disruption of cholesterol. It is worth noting that all four inhibitors negatively impacted the membrane binding capability of S4. Moreover, overdose treatment of endocytosis inhibitors caused cells to be unhealthy. For example, 4 mg*/*ml treatment of Poly-I and 20 µM treatment of Pitstop-2 stressed cells and disrupted the membrane binding of S4. Overall, these results show that the endocytic pathway of membrane-bound DNs is similar to bare DNs, which is an interdependent process that involves scavenger receptors, clathrin- and caveolin-mediated endocytosis.

**Figure 5:**
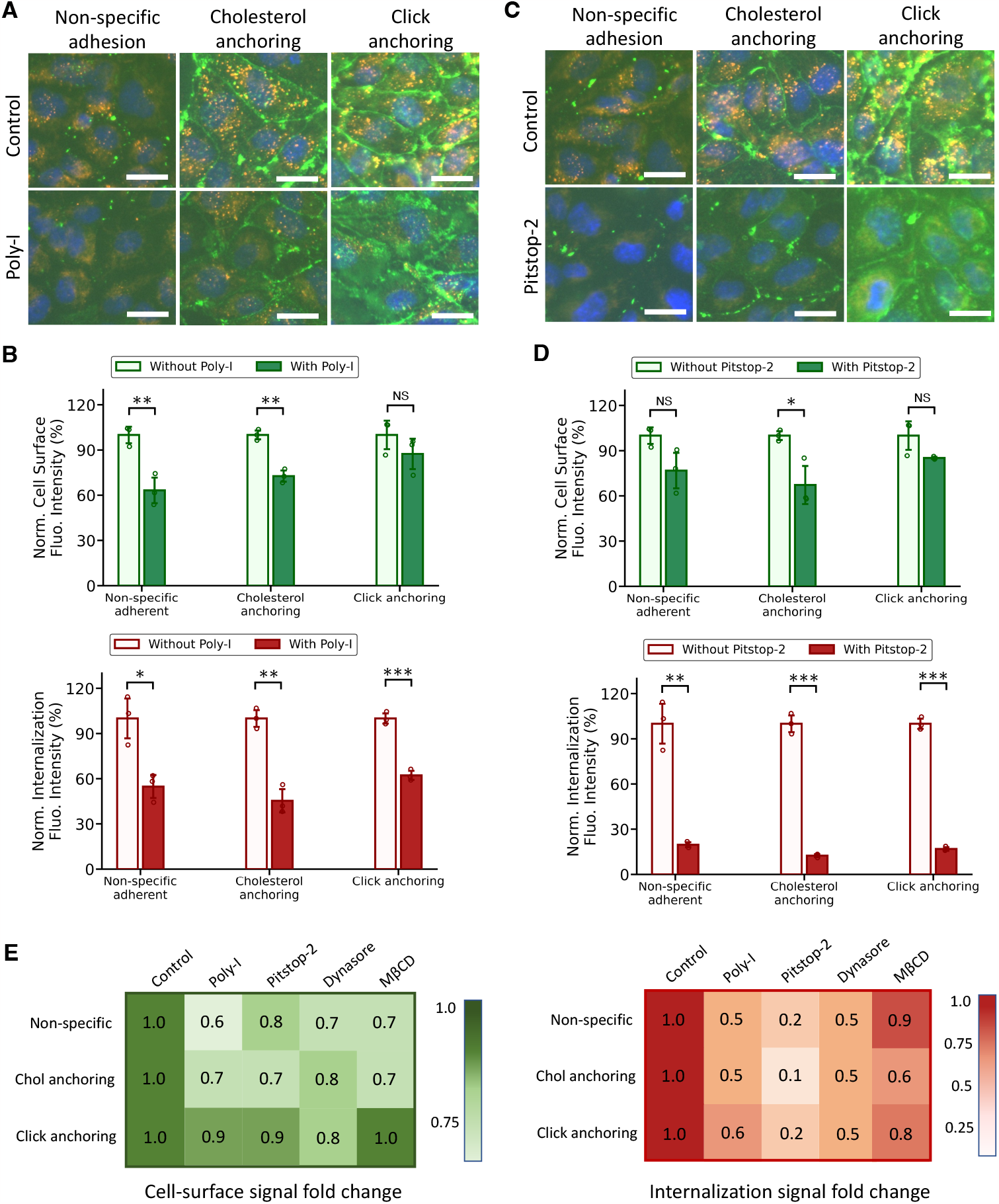
Endocytosis inhibition study of membrane-bound nanpspheres with 4 edge-decorated overhangs. (A) Fluorescence images of cells treated with and without the inhibitor of scavenger receptors, Poly-I. Cells were incubated 30 min at 37°C with 400 µg*/*ml of Poly-I, followed by fixation and staining. (B) Quantification of membrane-bound and internalization signal intensities for cells incubated with and without Poly-I. (C) Fluorescence images of cells treated with and without the inhibitor of clathrin-mediated endocytosis, Pitstop-2. Cells were incubated 30 min at 37°C with 10 µM of Pitstop-2, followed by fixation and staining. (D) Quantification of membrane-bound and internalization signal intensities for cells incubated with and without Pitstop-2. (E) Heat maps of cell-surface and internalization signal intensities fold change with endocytosis inhibitors. In each membrane binding method, data were normalized to the control group. In fluorescence images, membrane-bound S4 were stained with streptavidin-AF488 (green) and internalized DNs were stained with streptavidin-AF647 (red). Cell nuclei were stained with Hoechst (blue). Data were presented as means ± s.d. in (B) and (D) with n=3. \**P ≤*0.05, \*\**P ≤*0.01, \*\*\**P≤* 0.001. All scale bars: 10 µm.

## Conclusions

To summarize, we comprehensively investigated the enhancement effect of the membrane binding on the uptake of DNs. Three membrane binding strategies, non-specific adhesion, cholesterol membrane anchoring (one-step and two-step) and click glycocalyx anchoring (two-step), were studied. Three shapes of DNs, nanospheres, nanorods and nanotiles, with various decorations of ssDNA overhangs, were synthesized and their membrane binding capability and subsequent uptake were assessed. For two-step cholesterol anchoring and click anchoring methods, we found that the cellular uptake of DNs was significantly enhanced up to 8-fold. The effect of geometry on cellular uptake was also demonstrated: nanospheres, in general, were the most preferably internalized among all structures. But the placement of ssDNA overhangs may compensate for deficiencies in geometry. For example, edge-decorated nanotiles were found to outperform the nanospheres and were the most internalized in all 9 DNs. Furthermore, we demonstrate that although having a lower efficiency of membrane binding and uptake, one-step cholesterol binding approach provides simpler route for robust cell surface labeling and uptake. Our study on the endocytic pathways of membrane-bound DNs indicates that they have similar pathways as bare DNs. Thus same inhibitors may still be employed to effectively block their uptake. In general, these results show that our membrane- and glycocalyx-binding approach might be applied to not only a broader category of DNA nanostructures, but also various membrane binding strategies, providing insights into leveraging DNs as cellular delivery vehicles for biosensing, drug delivery, and gene therapeutics.

## Acknowledgement

This work was supported by grants from the Air Force Office of Science Research (Award no. FA9550-18-1-0199), the National Science Foundation (Award no. 2113301 to CMMI) and a 2022 Dowd Fellowship. The AFM imaging studies were performed using instrumentation funded by Defense University Research Instrumentation Program (Award no. FA9550-22-1-0147). The authors thank the Department of Biomedical Engineering, Molecular Biosensor and Imaging Center (MBIC) at Carnegie Mellon University, and Dr. Lydia A. Perkins for assistance on confocal microscope imaging.

