## Supplementary Information for "Membrane and glycocalyx tethering of DNA nanostructures for enhanced uptake"

### Membrane binding-enhanced cellular uptake of DNA nanostructures

∗ indicates equal author contribution

**List of Supplementary Information:**

Supplementary Figure 1. Fluorescence image processing by using morphological transformation and adaptive thresholding to reduce background noise.

Supplementary Figure 2. Quantification of cell-surface, internalization signals and internalization efficiency using cholesterol anchoring with absolute values.

Supplementary Figure 3. Quantification of cell-surface, internalization signals and internalization efficiency using click anchoring with absolute values.

Supplementary Figure 4. Time-dependent membrane binding and uptake of DNs.

Supplementary Figure 5. DN concentration-dependent membrane binding and uptake of DNs.

Supplementary Figure 6. Cholesterol anchor concentration-dependent membrane binding and uptake of DNs.

Supplementary Figure 7. Buffer-dependent membrane binding and uptake of DNs.

Supplementary Figure 8. Schematic illustrations of membrane wrapping of DNs.

Supplementary Figure 9. Endocytosis inhibition study by using dynasore to inhibit clathrin-mediated endocytosis.

Supplementary Figure 10. Endocytosis inhibition study by using MβCD to inhibit caveolin-mediated endocytosis.

Materials and Methods

Table S1. Calculation of membrane deformation energy needed for the spontaneous membrane wrapping of DNs.

Table S2. List of functional DNA oligos.

Table S3. List of DNA oligos for DNA nanospheres.

Table S4. List of DNA oligos for DNA nanorods.

Table S5. List of DNA oligos for DNA nanotiles.

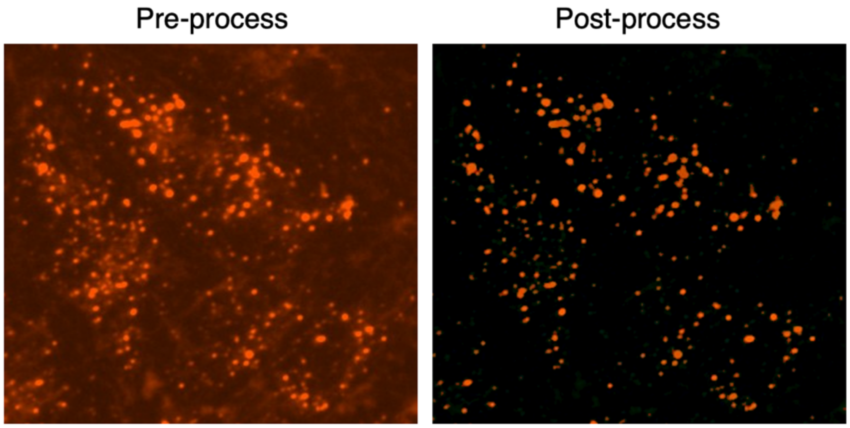

Supplementary Figure 1. Fluorescence image processing by using morphological transformation and adaptive thresholding to reduce background noise.[1,2]

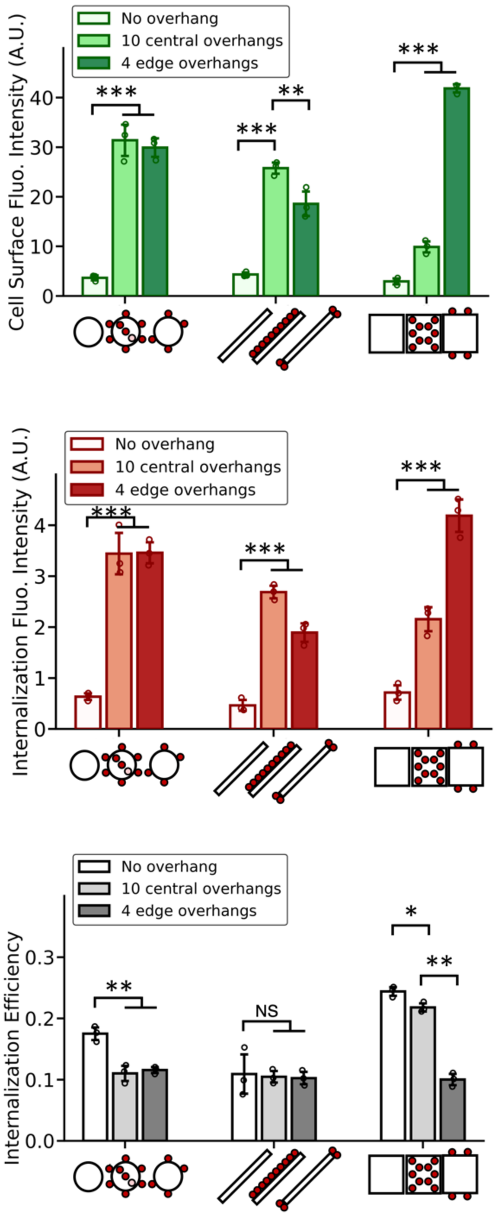

Supplementary Figure 2. Quantification of cell-surface, internalization signals and internalization efficiency using cholesterol anchoring by absolute values. Data were presented as means ± s.d. with n=3. *P ≤ 0.05, **P ≤ 0.01, ***P ≤ 0.001.

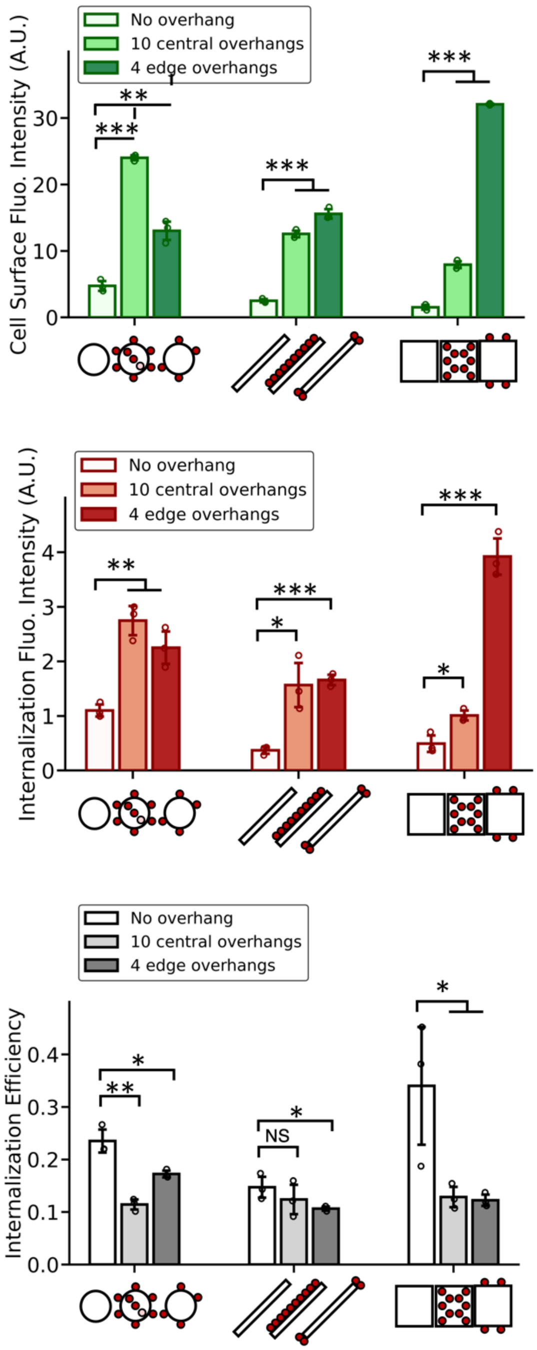

Supplementary Figure 3. Quantification of cell-surface, internalization signals and internalization efficiency using click anchoring by absolute values. Data were presented as means ± s.d. with n=3. *P ≤ 0.05, **P ≤ 0.01, ***P ≤ 0.001.

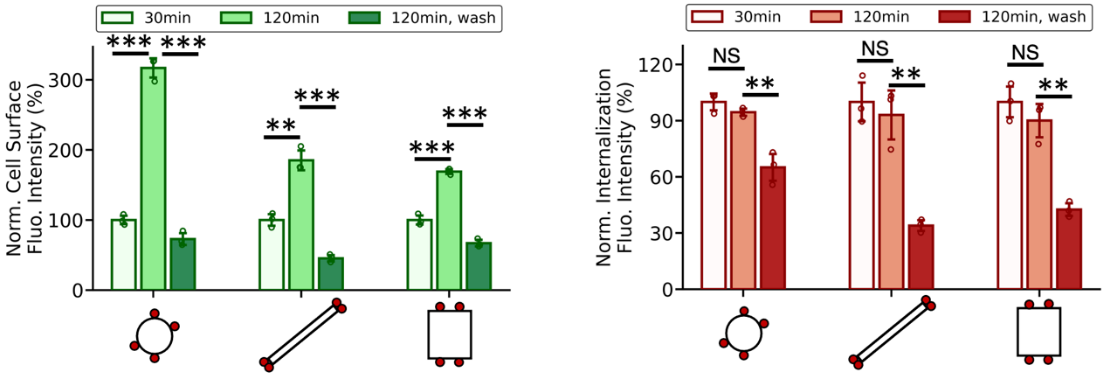

Supplementary Figure 4. Time-dependent membrane binding and uptake of DNs. Cells were incubated with DNs for 30 min, 120 min and 30 min followed by washing and continuing incubating for 90 min. All data were normalized to 30 min incubation group. **P ≤ 0.01, ***P ≤ 0.001.

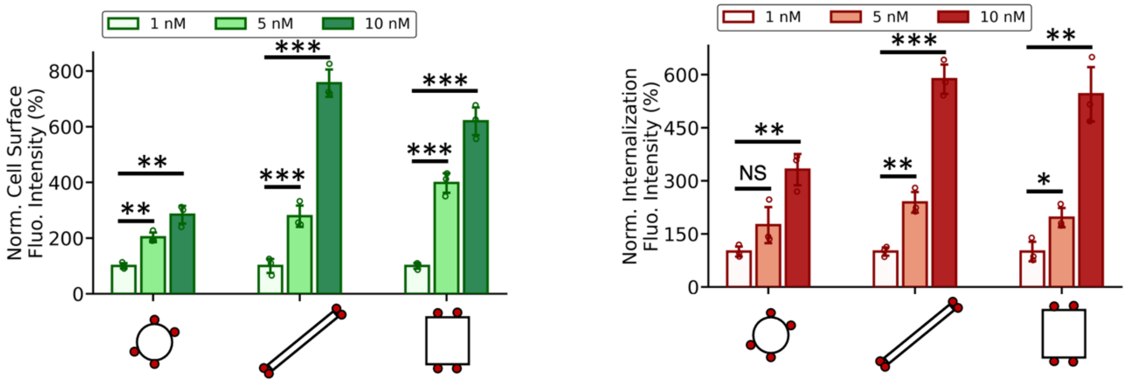

Supplementary Figure 5. DN concentration-dependent membrane binding and uptake of DNs. The concentration of DNs were changed from 1 nM, 5 nM to 10 nM. All data were normalized to 1 nM group. *P ≤ 0.05, **P ≤ 0.01, ***P ≤ 0.001.

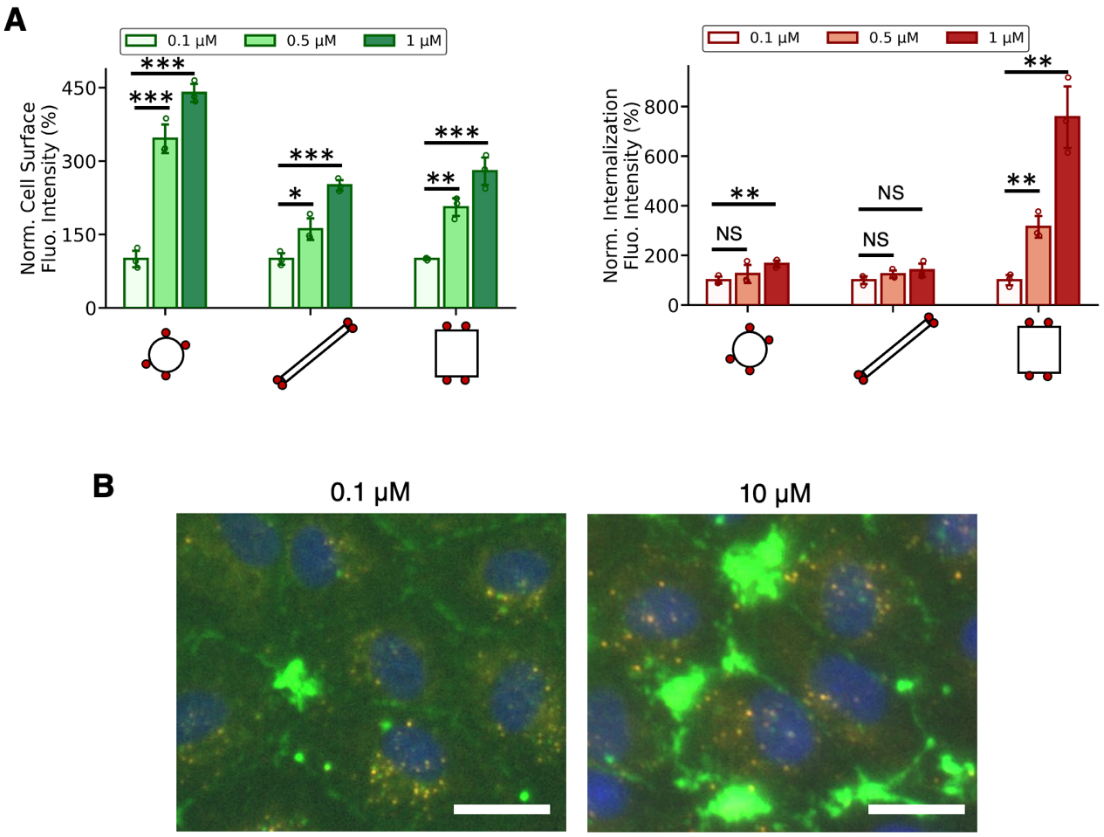

Supplementary Figure 6. (A) Cholesterol anchor concentration-dependent membrane binding and uptake of DNs. The concentration of cholesterol anchors were changed from 0.1 μM, 0.5 μM to 1 μM. All data were normalized to 0.1 nM group. (B) Aggregations triggered by a high concentration of cholesterol anchors. Scale bars: 10 μm. *P ≤ 0.05, **P ≤ 0.01, ***P ≤ 0.001.

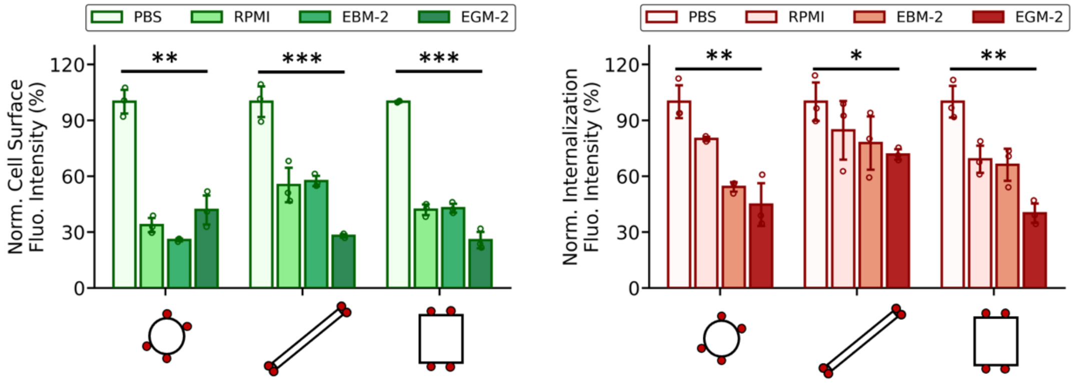

Supplementary Figure 7. Buffer-dependent membrane binding and uptake of DNs. Cells were incubated with DNs in PBS, RPMI, EBM-2 and EGM-2 for 30 min. All data were normalized to PBS buffer group. *P ≤ 0.05, **P ≤ 0.01, ***P ≤ 0.001.

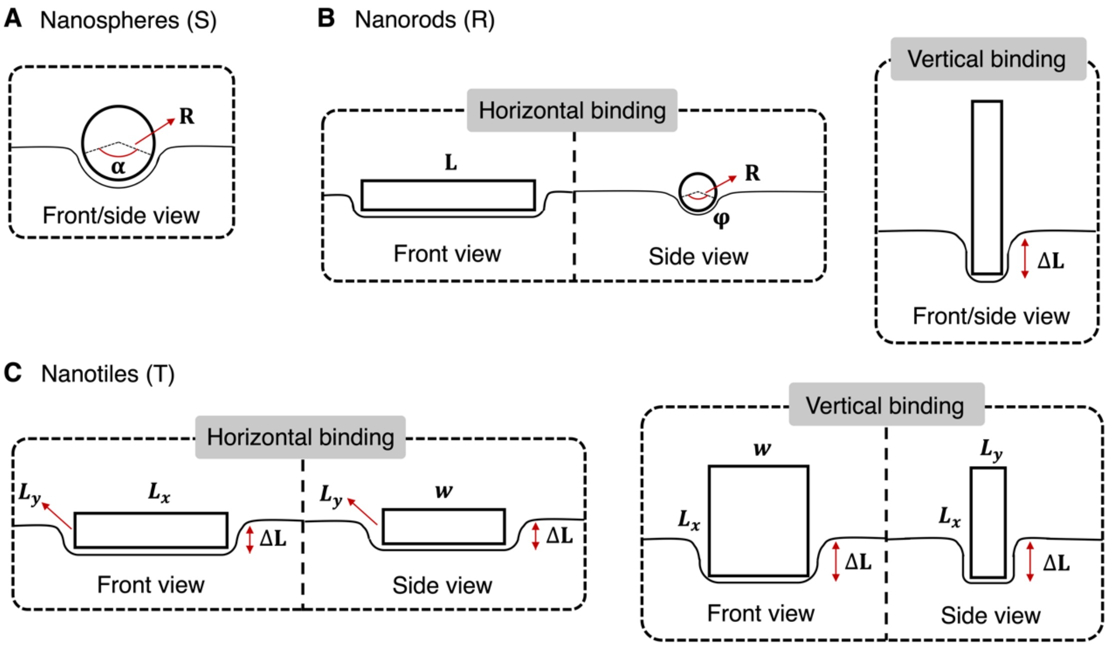

Supplementary Figure 8. Schematic illustrations of membrane wrapping of DNs, including (A) DNA nanospheres, (B) nanorods and (C) nanotiles. For each DN, two approaches of membrane wrapping were analyzed, including DN horizontal binding and vertical binding to membranes. $\varphi$ and $\Delta L$ denote the angle and length of the DN that had been wrapped by the lipid membranes, respectively. The diameter of nanospheres is R. The diameter of the nanorod is $R$ and its length is $L$. The dimension of the nanotile is $L_x * L_y * w$.

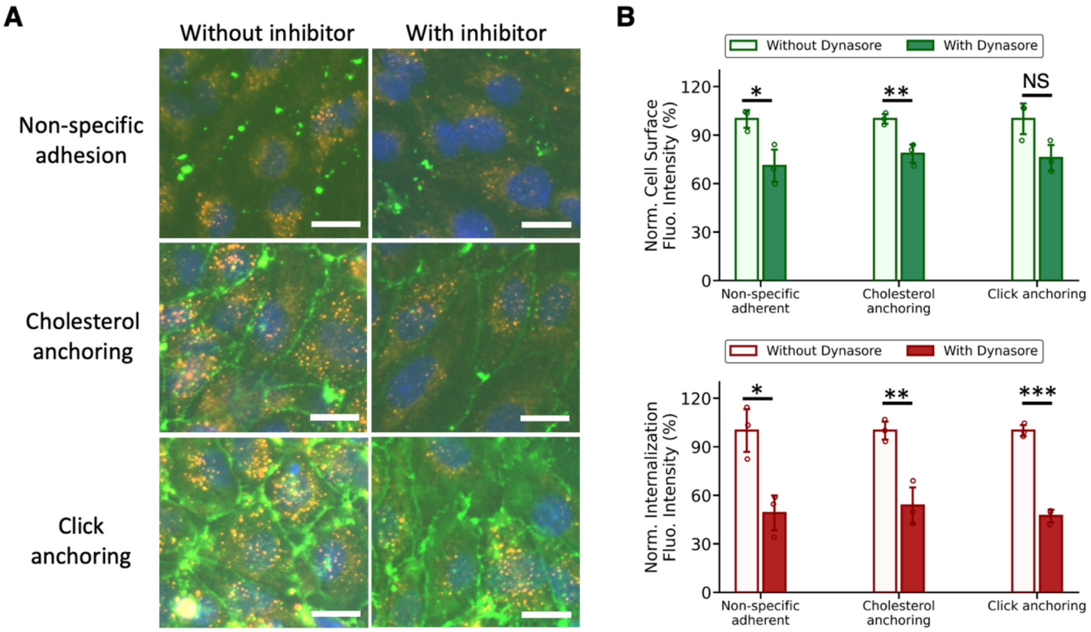

Supplementary Figure 9. Endocytosis inhibition study by using Dynasore to inhibit clathrin-mediated endocytosis. (A) Fluorescence images after incubating cells with and without Dynasore. Cells were incubated 30 min at 37°C with 120 µM of Dynasore, followed by fixation and staining. All scale bars: 10 µm. (B) Quantification of cell-surface and internalization signal intensities for cells incubated with and without Dynasore. $^*P \leq 0.05$, $^{**}P \leq 0.01$, $^{***}P \leq 0.001$.

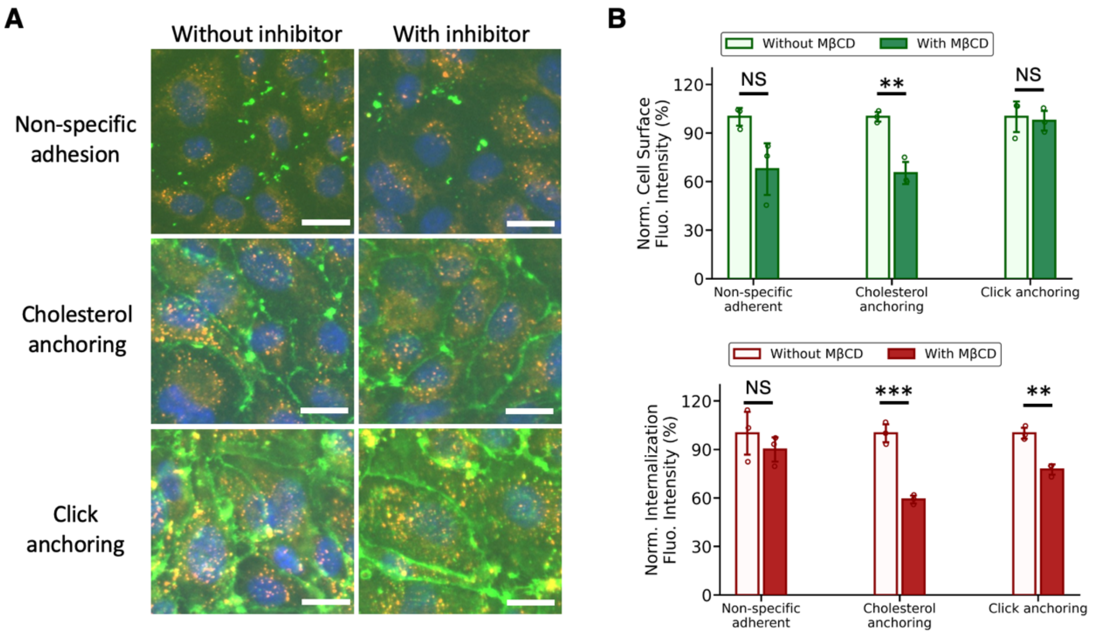

Supplementary Figure 10. Endocytosis inhibition study by using MβCD to inhibit caveolin-mediated endocytosis. (A) Fluorescence images after incubating cells with and without MβCD. Cells were incubated 30 min at 37°C with 300 nM of MβCD, followed by fixation and staining. All scale bars: 10 µm. (B) Quantification of cell-surface and internalization signal intensities for cells incubated with and without MβCD. **P ≤ 0.01, ***P ≤ 0.001.

#### Materials and Methods

##### Materials, reagents and equipment

DNA oligos, cholesterol-modified ssDNA, amine-modified ssDNA and biotin-modified ssDNA (Supplementary Table. 2, 3, 4 and 5) were purchased from Integrated DNA Technologies (Coralville, IA). DBCO-cy5 and DBCO-Sulfo-NHS-Ester, Sodium Chloride and Paraformaldehyde and PEG 8000 were purchased from Sigma-Aldrich. Streptavidin-Alexa Fluor 488 and 647 conjugates were purchased from Invitrogen. Roswell Park Memorial Institute (RPMI) 1640 Medium, Dulbecco's Phosphate-Buffered Saline (DPBS) was purchased from Corning. AFM tips were purchased form NanoAndMore (OMCL-AC160TS). SYBR Safe DNA gel stain was purchased from Invitrogen. Human umbilical vein endothelial cells, Endothelial Cell Growth Medium-2 BulletKit was purchased from Lonza.

##### Synthesis and purification of biotinylated DNA nanostructures

Synthesis and purification of biotinylated DNs All DNs were folded from M13mp18 scaffold (Bayou Biolabs) together with staple strands (Supplementary Table. 3, 4 and 5) through custom annealing ramps. Please refer to our previous studies for Cadnano desgin.[3,4] For annealing, 20 nM of the scaffold and 100 nM of the staples were mixed in TAE buffer with 12.5 mM $MgCl_2$ (nanorods and nanotiles) and with 20 mM $MgCl_2$ for nanospheres. The biotin-labeled strands were added pre-annealing with the same concentration as other staples. Cholesterol-conjugated DNs used 3x excess of cholesterol-conjugated staples. The mixture for synthesizing nanotiles was heated to 90°C for 5 min and gradually cooled down to 4°C within ~4 hours. Specifically, 90°C-70°C, 0.1°C per 6 seconds; 70°C-45°C, 0.1°C per 30 seconds; 45°C-30°C, 0.1°C per 6 seconds; 30°C-4°C, 0.1°C per 3 seconds. The mixture for synthesizing nanospheres and nanorods was heated to 80°C for 5 min and gradually cooled down to 4°C within ~45 hours. Specifically, 80°C-65°C, 0.1°C per 24 seconds; 65°C-24°C, 0.1°C per 378 seconds; 24°C-4°C, 0.1°C per 18 seconds.

After annealing, the solutions were collected and precipitated by centrifugation at 10500 g for 25 min in a buffer containing 7.5% PEG 8000, 10 mM $MgCl_2$, 255 mM NaCl, 22.5 mM Tris, 10 mM acetic acid, and 1 mM EDTA. The supernatant was removed and the remaining was resuspended to 5 nM using DPBS buffer with 12.5 mM $MgCl_2$. DNs were then administrated to cells within 30 min after purification.

**Characterizations of DNs**

For agarose gel electrophoresis, 10 µL of 10 nM purified DNs samples were first incubated in DPBS buffer for 30 min at 37°C C, and then analyzed by electrophoresis in 2% agarose gel in TBE buffer with 12.5 mM MgCl2 at 90 V and room temperature (RT) for 1 hour. The gels were stained with 1× SYBR Safe DNA gel stain and imaged with Biorad ChemiDoc Imaging System.

For atomic force microscopy (AFM) characterization, 10 µL of purified DNs samples (1-2 nM in concentration) in TAE with 12.5 mM $MgCl_2$ was deposited onto a freshly cleaved mica surface, incubated at RT in a humid chamber for 5 min. The sample was washed three times with 20 µL DI water and thoroughly blow-dried with nitrogen after each washing. AFM scans were performed using NX10 AFM system with OMCL-AC160TS tips pre-loaded onto a wafer (Park Systems Corp.), in non-contact mode (NCM).

**Synthesis of DBCO-ssDNA**

The $NH_2$-ssDNA oligos were incubated overnight in DPBS with DBCO-sulfo-n-hydroxysuccinimidyl ester (DBCO-Sulfo-NHS-Ester) at a 1:10 molar ratio under agitation at RT. 30 min After incubation, the reaction mixture was dialyzed five times against DPBS using Amicon Ultra Centrifugal filters (molecular weight cut-off, 3 kDa) to remove unconjugated DBCO-Sulfo-NHS-Ester. The above-described conjugation and purification procedures were repeated twice in total to increase the conjugation efficiency. The final concentration of DBCO-ssDNA was adjusted to 500 µL with DPBS.

**Cell culture**

Human umbilical vein endothelial cells (HUVECs) were cultured in Endothelial Cell Growth Medium-2 (EGM-2) at 37°C C with 5% CO2. HUVECs of passage 3-5 were plated into 96-well culture plates at a density of 50,000 cells per well, one day prior to the experiment.

**Two-step cholesterol membrane anchoring**

First, cholesterol-ssDNA was diluted to 0.5 µM in DPBS and administered to cells followed by one-hour incubation at 37°C C. 5 µM of DNs bearing complementary ssDNA overhangs were then introduced to cells followed by 30 min incubation. Cells were then fixed and the on-membrane DNs and internalized DNs were visualized via dual-color streptavidin staining.

**Two-step click glycocalyx anchoring**

First, azide ligands were labeled on cell-surface glycocalyx through metabolic labeling. Specifically, azido monosaccharide, N-azidoacetylmannosamine-tetraacylated (Ac4ManNAz, stock in DMSO) was diluted in culture medium to a final concentration of 50 µM and administered to cells for two days prior to the experiment. Next, cells were incubated with DBCO-ssDNA at 50 µM in DPBS for 1 hour at 37°C to allow DBCO-azide click conjugation. 5 µM of DNs bearing complementary ssDNA overhangs were then introduced to cells followed by 30 min incubation. Cells were then fixed and the on-membrane DNs and internalized DNs were visualized via dual-color streptavidin staining.

**One-step cholesterol membrane anchoring**

5 µM of DNs with direct conjugation of cholesterol tags were introduced to cells followed by 30 min incubation at 37°C. Cells were then fixed and the on-membrane DNs and internalized DNs were visualized via dual-color streptavidin staining.

**Dual-color streptavidin staining of membrane-bound and internalized DNs**

After incubating with DNs, cells were fixed with PFA and first stained with streptavidin-Alexa Fluor 488 in 1% BSA in DPBS for 30 min to visualize membrane-bound DNs. Cells were then permeabilized with 0.5% Triton-X for 15 minutes at RT. The internalized DNs were then visualized via incubation of the permeabilized cells with the streptavidin-Alexa Fluor 647 in 1% BSA in DPBS for 30 min at RT.

**Quantitative image analysis of internalized DNs**

To minimize background noise caused by non-specific cell internalization of fluorophores, we performed image analysis to only count for internalized DN signals.[5] Specifically, internalized signals were identified by constructing structuring elements and performing morphological transformations (Fig. S2).[1,2] The intensities of post-processed images were then calculated by summing up the intensity of all pixels. For on-membrane signals, the intensity of each image was directly computed without processing.

**Adhesion-driven envelopment of DNs**

To study if the internalization of membrane-bound DNs is a spontaneous membrane wrapping process driven by adhesion, we calculated the adhesion energy provided by the insertion of cholesterol tags into lipid membranes, and the energy needed for a membrane to wrap, engulf, and finally internalize a DN. Here lipid membranes were modeled as simple elastic continuum. The adhesion energy provided by cholesterol insertion is $E_{ahesion} = n(x * k_B T)$. $n$ is the number of cholesterol tags inserted into the membrane and $x * k_B T$ is the adhesion gained from the insertion of one cholesterol. A membrane's bending energy can be described by Helfrich expression.[6] For nanorods and nanotiles, we considered two possible orientations when they interact with cell membranes, vertically and horizontally (Supplementary Figure 8). We found that for all DNs, the adhesion energy was substantially smaller than what would be needed to drive passive uptake (Table. S1). Here we used nanospheres as an example. Suggested by previous theoretical and experimental calculations, we took the bending modulus of the cellular membrane $\kappa$ to be $25k_B T$ and $x = 13$.[7,8] The radius of the nanosphere is 54 nm. Then the membrane bending energy is $E_{bending} = \frac{1}{2}\kappa(\frac{1}{R} + \frac{1}{R})^2 A_{sphere} = 8\pi\kappa \approx 628k_B T$. For spontaneous wrapping to happen, $E_{bending}$ must be smaller or equal to $E_{ahesion}$. We then got $n \geq 48$ for nanospheres, meaning we need at least 48 cholesterol tags on a nanosphere for spontaneous wrapping, which is a very big number and we don't have these many cholesterol tags on our nanospheres. Thus active endocytosis must be involved in the internalization of membrane-bound DNs.

**Inhibition of endocytic pathways study**

Inhibitors were diluted using EBM-2 to a certain concentration. Specifically, Poly-I was 400 µg/ml. Pitstop-2 was 10 µM. Dynasore was 120 µM. MβCD was 300 nM. Cells were first incubated with inhibitors for 30 min at 37°C. Then cells were washed and administrated with cholesterol anchors for two-step cholesterol anchoring, or DBCO-ssDNA for two-step click anchoring for 30 min at 37°C. For control group, EBM-2 medium was added for incubation. After incubation, cells were washed again and incubated with DNs for another

30 min at 37°C, followed by washing, fixation and staining. Fluorescence microscopy images of cells were taken and assessed for quantification.

**Statistics and Reproducibility**

Quantitative data were displayed as means ± s.d. Statistical significances were determined using one-way analysis of variance (ANOVA) with post-hoc Tukey's Test. All cell experiments were repeated independently at least three times.

Table S1. Calculation of energy needed for spontaneous wrapping of DNs. See Supplementary Figure. 9 for schematic illustrations. The adhesion energy provided by the interaction between DNs and cell membranes (the insertion of cholesterol into lipid membranes) is $n(k_B T)$. $n$ is the number of cholesterol tags inserted into the membrane. $\kappa$ is the bending modulus of the cellular membrane. In the nanotiles, the curvature radius for membrane bending is related to the tile thickness $L_y$.

| DNs | | Membrane bending energy to wrap the DN |
|---|---|---|
| Nanosphere | Horizontal | $\frac{1}{2}\kappa(\frac{1}{R}+\frac{1}{R})^2 A_{sphere} = 8\pi\kappa$ <br> R is the radius of sphere. |
| | Vertical | |
| Nanorod | Horizontal | $\frac{1}{2}\kappa(\frac{1}{R})^2 A_{rod,horizontal} = \frac{\kappa}{2R}\varphi L$ <br> R is the radius of the rod. φ the portion of the rod that has been engulfed. |
| | Vertical | $\frac{1}{2}\kappa(\frac{1}{R})^2 A_{rod,vertical} = \frac{\pi\kappa\Delta L}{R}$ <br> R is the radius of the rod. ΔL is the portion of the rod that has been engulfed. |
| Nanotile | Horizontal | $\frac{1}{2}\kappa(\frac{1}{R})^2 A_{tile,horizontal} = \frac{1}{2}\kappa\left(\frac{1}{R}\right)^2 2(L_x + w)\Delta L$ <br> R is the membrane curvature near the edge of tile. $\Delta L$ is the portion of the tile that has been engulfed. |
| | Vertical | $\frac{1}{2}\kappa(\frac{1}{R})^2 A_{tile,vertical} = \frac{1}{2}\kappa\left(\frac{1}{R}\right)^2 2(L_y + w)\Delta L$ <br> R is the membrane curvature near the edge of tile. $\Delta L$ is the portion of the tile that has been engulfed. |

Table S2. List of functional DNA oligos.

| Name of oligos | Sequence |
|---|---|
| ssDNA on DNs for two-step membrane binding (both cholesterol membrane anchoring and click glycocalyx anchoring) | 5'/TT CAGTCAGTCAGTCAGTCAGT/3' |
| ssDNA on DNs for attaching biotin | 5'/TT GAGAGCAGACCTGGAACTCG/3' |
| Complementary ssDNA conjugated with biotin | 5'/BiosG/TT CGAGTTCCAGGTCTGCTCTC/3' |
| Cholesterol-ssDNA anchors for immobilizing cell membranes in two-step cholesterol anchoring | 5'/Chol-TEG/ACTGACTGACTGACTGACTG/3' |
| 5'$NH_2$-ssDNA for synthesizing DBCO-ssDNA anchors in click anchoring | 5'/AmMC6/TT ACTGACTGACTGACTGACTG/3' |
| Cholesterol sequence for one-step cholesterol membrane binding | 5'/Chol-TEG/AAACTGACTGACTGACTG/3' |
| Shielding adjacent ssDNA on DNs for one-step cholesterol membrane binding | 5'/TT CAGTCAGTCAGTCAGTTTCCATCA/3' |

Table S3. List of DNA oligos for DNA nanospheres

| | | | |
|---|---|---|---|
| CCACCCTCAGAGCCACCACCCTCATAGCTATCTTACCGAAGCCCT | | | |
| AGCCACCACCGGAACCGCCTCCCACTATATGTAAATGCTGATGCAAATCC | | | |
| TTATTAGCGTTTGCCATCTTTTCATGTTAGCAAACGTAGAAAAT | | | |
| CGTCAGACTGTAGCGCGTTTTCAATCATATGCGTTATACAAATTCTTACC | | | |
| GATAGCAGCACCGTAATCAGTAGAAAGGTGAATTATCACCGTCACCG | | | |
| ACCATTACCATTAGCAAGGCCATGTTCAGCTAATGCAGAACGCGC | | | |
| ACTTGAGCCATTTGGGAATTAGATTTCATCGTAGGAATCATTAC | | yes (center) | yes (edge) |
| AATATTGACGGAAATTATTCATTTATAAAAGAAACGCAAAGACACCA | | | |
| AGCGCCAAAGACAAAAGGGCGACATTCGAGCGTCTTTCCAGAGCCTAATT | | | |
| CGGAATAAGTTTATTTTGTCACAGGAATACCCAAAAGAACTGGCATG | | | |
| ACATACATAAAGGTGGCAACACGACAGAATCAAGTTTGCCTTTAG | | yes (center) | |
| ATTAAGACTCCTTATTACGCAGTAGCCGAACAAAGTTACCAGAAGGA | | | |
| AACCGAGGAAACGCAATAATAACAATTGAGTTAAGCCCAATAATAAG | | | |
| TTTTAAGAAAAGTAAGCAGATATAATCAAAATCACCGGAACCAG | | | |
| AGCAAGAAACAATGAAATAGCAATACCGTTCCAGTAAGCGTCATACATGG | | yes (center) | |
| ATATCAGAGAGATAACCCACAAGCAAAAATGAAAATAGCAGCCTTTA | | | |
| ATTAACTGAACACCCTGAACAGATTAGCGGGGTTTTGCTCAGTAC | | | |
| CAGAGAGAATAACATAAAAACAGATATTATTTATCCCAATCCAAATA | | | |
| AGAAACGATTTTTTGTTTAACGTATCAATAGAAAATTCATATGGTTTACC | | | |
| TGCCAGTTACAAAATAAACAGCCAGGTTTAGTACCGCCACCCTC | | | |
| ATCCTGAATCTTACCAACGCTAACAGATATAGAAGGCTTATCCGGTA | | | |
| TGAAGCCTTAAATCAAGATTAGTTGCAGTTTTGTCGTCTTTCCAGACGTT | | | |
| TTCTAAGAACGCGAGGCGTTTTAATTAAACCAAGTACCGCACTCATC | | | |
| CGCGCCCAATAGCAAGCAAATCAACCGATTGAGGGAGGGAAGGTA | | | |
| GAGAACAAGCAAGCCGTTTTTATAACCAATCAATAATCGGCTGTCTT | | | |
| TCCTTATCATTCCAAGAACGGGTATAGTTGCGCCGACAATGACAACAACC | | | |
| ATCCTAATTTACGAGCATGTAGAGCCAGCAAAATCACCAGTAGC | | | |
| CTGTTTATCAACAATAGATAAGTCCATTAAACGGGTAAAATACGTAATGC | | | |
| TCCAGACGACGACAATAAACAACGGCTTAATTGAGAATCGCCATATT | | | |
| GTAATAAGAGAATATAAAGTAACCTGTCGTGCCAGCTGCATT | | | |
| TAACAACGCCAACATGTAATTTATAAGAATAAACACCGGAATCATAA | | | |
| AGTATAAAGCCAACGCTCAACAGTAGGGAAACGTCACCAATGAAACCATC | | | |
| TTACTAGAAAAAGCCTGTTTAGTACTTTTTCAAATATATTTTAGTTA | yes | | |
| TGTGATAAATAAGGCGTTAAAGGAGAGGCGGTTTGCGTATTGGGC | | | |
| ATTTCATCTTCTGACCTAAATTTGTCTGAGAGACTACCTTTTTAACC | | | |
| AATCGCAAGACAAAGAACGCGAGAAATCGGCATTTTCGGTCATAGCCCCC | yes | | |
| TCCGGCTTAGGTTGGGTTATATATTAATTTTCCCTTAGAATCCTTGA | | | |
| GTGAATTTATCAAAATCATAGAGAGTTGCAGCAAGCGGTCCACGC | | | |
| AAACATAGCGATAGCTTAGATTACATTTAACAATTTCATTTGAATTA | | | |
| TTGCTTCTGTAAATCGTCGCTATTAATCAGAGCCGCCACCCTCAGAACCG | | | |
| CCTTTTTTAATGGAAACAGTACAAGCAAAAGAAGATGATGAAACAAA | | | |
| CATCAAGAAAACAAAATTAATTAAAAGAATAGCCCGAGATAGGG | yes | | |
| GAATTATTCATTTCAATTACCTGATTGCGTAGATTTTCAGGTTTAAC | | | |
| CGGATTCGCCTGATTGCTTTGAATACGGGAGCCCCCGATTTAGAGCTTGA | | | |
| GTCAGATGAATATACAGTAACAGTCCTGATTGTTTGGATTATACTTC | yes | | |
| ACGTAAAACAGAAATAAAGAAAGGTTGAGGCAGGTCAGACGA | | | |
| TGAATAATGGAAGGGTTAGAACCAACAGTTAATGCCCCCTGCCT | | | |
| ATGATGGCAATTCATCAATATAACAAGTGTAGCGGTCACGCTGCGCGTAA | yes | | |
| GAAACCACCAGAAGGAGCGGAATGAGGATTTAGAAGTATTAGACTTT | | yes (center) | yes (edge) |
| ATTAATTTTAAAAGTTTGAGTTGATAGCCCTAAAACATCGCC | | | |
| ACAAACAATTCGACAACTCGTATTGGCAAATCAACAGTTGAAAGGAA | yes | | |
| TAGAGCCGTCAATAGATAATACATTTAAACATGAAAGTATTAAGAGGCTG | | | |
| TTGAGGAAGGTTATCTAAAATATTAGGAACCCATGTACCGTAACACTGAG | | | |
| CCCTCAATCAATATCTGGTCAGTACCAGCAGAAGATAAAACAGAGGT | yes | | |
| CAGCAAATGAAAAATCTAAAGCATCATGGCTCATTATACCAGTCAGGACG | | | |
| GAGGCGGTCAGTATTAACACCGCGGCACAGACAATATTTTTGAATGG | | | |
| ATTAAAAATACCGAACGAACCTAAATCCTTTGCCCGAACGTT | | | |
| CTATTAGTCTTTAATGCGCGAACAAAGGGATTTTAGACAGGAACGGTACG | | | |
| GACCTGAAAGCGTAAGAATACGTAGTTGAGATTTAGGAATACCA | | | |
| ACCAGTAATAAAAGGGACATTCTACAATATTACCGCCAGCCATTGCA | | | |
| CGTCTGAAATGGATTATTTACGCGTCCAATACTGCGGAATCG | | | |
| ACAGGAAAAACGCTCATGGAAATTTGCGGATGGCTTAGAGCTTAATTGCT | | | |
| CGGCCTTGCTGGTAATATCCAGAATCAGTGAGGCCACCGAGTAAAAG | | | |
| GATTAGTAATAACATCACTTGGTTTTTTGGGGTCGAGGTGCCGTA | | | |
| AGTCTGTCCATCACGCAAATTAATGCTTTCCTCGTTAGAATCAGAGC | | | |
| CCAGAATCCTGAGAAGTGTTTTTATAGGCCAACAGAGATAGAACCCTTCT | | | |

| | | | |
|---|---|---|---|
| GGGAGCTAAACAGGAGGCCGATTCCGCTACAGGGCGCGTACTATGGT | | | |
| TGCTTTGACGAGCACGTATAACGGAGAAAGGAAGGGAAGAAAGCGAA | | | |
| CCACCACACCCGCCGCGCTTAATGCGAACATTATCATTTTGCGGAACAAA | | | |
| AGGAGCGGGCGCTAGGGCGCTGGTACCTTTTACATCGGGAGAAACAATAA | | | |
| CGGGGAAAGCCGGCGAACGTGGCCCGTTGTAGCAATACTTCTTT | | yes (center) | yes (edge) |
| AAGCACTAAATCGGAACCCTAAAAACAAGAGTCCACTATTAAAGAAC | | | |
| ACTACGTGAACCATCACCCAAATCAATTTAAATATGCAACTAAAGTACGG | | | |
| GTGGACTCCAACGTCAAAGGGCGTCCTGTTTGATGGTGGTTCCGAAA | | | |
| TTGAGTGTTGTTCCAGTTTGGCAAGTTACAAAATCGCGCAGAGGC | | | |
| TCGGCAAAATCCCTTATAAATCAAGACGCTGAGAAGAGTCAATA | | | |
| TGGTTTGCCCCAGCAGGCGAAAATAGATACATTTCGCAAATGGTCAATAA | | | |
| GCCCTTCACCGCCTGGCCCTGAGAATGGTTTGAAATACCGACCG | | | |
| GCCAGGGTGGTTTTTCTTTTCACACTCACATTAATTGCGTTGCGCTC | yes | | |
| AATGAATCGGCCAACGCGCGGGGCAGAGGCATTTTCGAGCCA | | yes (center) | yes (edge) |
| ACTGCCCGCTTTCCAGTCGGGAACTGATAAATTGTGTCGAAATCCGCGAC | | | |
| TGGGGTGCCTAATGAGTGAGCTAGCGCGAGCTGAAAAGGTGGCATCA | yes | | |
| ACAATTCCACACAACATACGACAAAAAGATTAAGAGGAAGCCCGA | | yes (center) | |
| ATTCTACTAATAGTAGTAGCATTAACAGTTGATTCCCAATTCTGCGA | | | |
| CCTGTTTAGCTATATTTTCATTTGGGCAGTGAGACGGGCAACAGCTGATT | yes | | |
| ACGAGTAGATTTAGTTTGACCATAAAAACCGTCTATCAGGGCGATGGCCC | | | |
| TGTCTGGAAGTTTCATTCCATATAGGATTAGAGAGTACCTTTAATTG | | | |
| GAATATAATGCTGTAGCTCAACATGTCCTGAGTAGAAGAACTCAAACTAT | | | |
| CTCCTTTTGATAAGAGGTCATTTATTCGAGCTTCAAAGCGAACCAGA | | | |
| CCGGAAGCAAACTCCAACAGGTCGTGTGAAATTGTTATCCGCTC | | | |
| AAGACTTCAAATATCGCGTTTTACCCTCAAATGCTTTAAACAGTTCA | yes | | |
| ATAGTCAGAAGCAAAGCGGATTGCATAGGCGCAGACGGTCAATCATAAGG | | | |
| GAAAACGAGAATGACCATAAATCAAGAAGTTTTGCCAGAGGGGGTAA | | | |
| TCATAAATATTCATTGAATCCACCTACATTTTGACGCTCAAT | | | |
| TAGTAAAATGTTTAGACTGGATATAACGCCAAAAGGAATTACGAGGC | | | |
| ACCAAAATAGCGAGAGGCTTTTGCAACTTCATCAAGAGTAATCTTGACAA | | yes (center) | |
| ATAGTAAGAGCAACACTATCATATAAAACGAACTAACGGAACAACAT | yes | | |
| CATTCAACTAATGCAGATACAATTGGCAGATTCACCAGTCACACG | | | |
| TATTACAGGTAGAAAGATTCATCCTGCAACAGTGCCACGCTGAGAGCCAG | | | |
| TTGGGAAGAAAAATCTACGTTAAAGAAACACCAGAACGAGTAGTAAA | | | |
| ACCTTATGCGATTTTAAGAACCCTGTAGCATTCCACAGACAGCCC | | | |
| TTGGGCTTGAGATGGTTTAATTTCTCCAAAAAAAAAGGCTCCAAA | | | |
| AGTGAATAAGGCTTGCCCTGACGACCCTCGTTTACCAGACGACGATAAAA | | | |
| GAACCGGATATTCATTACCCAAACACCCTCAGCAGCGAAAGACA | yes | | |
| ACCAGGCGCATAGGCTGGCTGACAAAAATCAGGTCTTTACCCTGACTATT | | | |
| GAACCGAACTGACCAACTTTGAAATTATACCAAGCGCGAAACAAAGT | | | |
| CTGCTCCATGTTACTTAGCCGGAACGGCCGGAAGCATAAAGTGTAAAGCC | | | |
| ACAACGGAGATTTGTATCATCGCAAAGAGGCAAAAGAATACACTAAA | yes | | |
| ACACTCATCTTTGACCCCCAGCGCGGCTACAGAGGCTTTGAGGACTA | | | |
| CACTACGAAGGCACCAACCTAAAACGCCGACAAAAGGTAAAGTAATTCTG | | | |
| AAGACTTTTTCATGAGGAAGTTTGGTCGCTGAGGCTTGCAGGGAGTT | | | |
| GCATCGGAACGAGGGTAGCAAAGAGGACAGATGAACGGTGTACAG | | | |
| AAAGGCCGCTTTTGCGGGATCGTTTTATCAGCTTGCTTTCGAGGTGA | | | |
| ATCGCCCACGCATAACCGATATATTCCCTGAACAAGAAAAATAATATCCC | | | |
| ATTTCTTAAACAGCTTGATACCGGGAACAACTAAAGGAATTGCGAAT | | | |
| AGGAGCCTTTAATTGTATCGGTCAACGTAACAAAGCTGCTCATTC | | | |
| AATAATTTTTTCACGTTGAAAATATTTTGCTAAACAACTTTCAACAG | | | |
| TTTCAGCGGAGTGAGAATAGAAAGCGAACCTCCCGACTTGCGGGAGGTTT | | | |
| AGTAAATGAATTTTCTGTATGGGCAACTTTAATCATTGTGAATT | | | |
| TCATAGTTAGCGTAACGATCTAAGCCACCCTCAGAGCCACCACCCTC | | | |
| TTTCGTCACCAGTACAAACTACAACGCCTTGCTGAACCTCAAATATCAAA | | | |
| ATTTTCAGGGATAGCAAGCCCAAGGGTTGATATAAGTATAGCCCGGA | | | |
| AGAACCGCCACCCTCAGAACCTATTTTGCACCCAGCTACAATTTT | | | |
| ATAGGTGTATCACCGTACTCAGGGGAAGCGCATTAGACGGGAGA | | | |
| CAGGCGGATAAGTGCCGTCGAGACTTTAGGAGCACTAACAACTAATAGAT | | | |
| AGACTCCTCAAGAGAAGGATTAGTAATAAGTTTTAACGGGGTCAGTG | | | |
| ATTTCGGAACCTATTATTCTGTATCATCATATTCCTGATTATCAG | | yes (center) | |
| CCTTGAGTAACAGTGCCCGTATAAAATAAATCCTCATTAAAGCCAGA | | | |
| CTTTTGATGATACAGGAGTGTACTGGAAGTCAGAGGGTAATTGAGCGCTA | | | |
| ATGGAAAGCGCAGTCTCTGAATTGAGCCGCCACCAGAACCACCACCA | | | |
| TTGGCCTTGATATTCACAAACTACCATATCAAAATTATTTGC | | | |
| GAGCCGCCGCCAGCATTGACAGGTAAATCAATATATGTGAGTGAATAACC | | yes (center) | |

Table S4. List of DNA oligos for DNA nanorod

| **DNA nanorod sequence** | biotin sites | 10 binding sites | 4 edge binding sites |
|---|---|---|---|
| TCAGGCTGCGCAACCTAGGGCGCTGGCAATCGTCTGAAATGG | | | |
| CATAACGCCAAAAGTTGCTAAACAACTTCCAATAGGAACCCA | | | |
| CCGCTTCTGGTGCCCCACACCCGCCGCGACAGGAAAAACGCT | | | |
| TATCGGCCTCAGGAATGGTTGCTTTGACTTGCTGGTAATATC | | | |
| CATCGTAACCGTGCGAATCAGAGCGGGAATAACATCACTTGC | | | |
| GATTGACCGTAATGTTAGACAGGAACGGTCACGCAAATTAAC | | | |
| TCAGTTGAGATTTAAAGGAACAACTAAACCACCCTCAGAGCC | | | |
| AACGAACTAACGGATGAAAATCTCCAAAGGTTTAGTACCGCC | | | |
| TATACCAGTCAGGAGTATCGGTTTATCAATATAAGTATAGCC | | | |
| ATCATTGTGAATTAAGCTTGATACCGATTTTTGCTCAGTACC | | | |
| CGAGTAGTAAATTGGCCCACGCATAACCAGAGGCTGAGACTC | | | |
| TCATTCAGTGAATAGAGTTAAAGGCCGCTGCCTATTTCGGAA | | | |
| AGAACCGGATATTCAAAGACAGCATCGGGTGCCTTGAGTAAC | | | |
| GGCGCATAGGCTGGTTGAGGACTAAAGAGATGATACAGGAGT | | | |
| TGACCAACTTTGAAGGGTAAAATACGTATCTCTGAATTTACC | | | |
| GCCGGAACGAGGCGCGAAAGAGGCAAAACAAACAAATAAATC | | | |
| GATAAATTGTGTCGCCCAGCGATTATACGAAGTATGTTGAGG | | | |
| TTGCGTATTGGGCGCTTTTCACCAGTGAAATAGATTAGAGCC | | | |
| TATCATAACCCTCGCGTCTTTCCAGACGGTACAAACTACAAC | | | |
| CAGCTGCATTAATGGCCTGGCCCTGAGATGAGGAAGGTTATC | | | |
| TTGCGCTCACTGCCTGCCCCAGCAGGCGATCAATATCTGGTC | | | |
| GCCTGGGGTGCCTATCGGCAAAATCCCTTCTAAAGCATCACC | | | |
| ACAATTCCACACAAGGTTGAGTGTTGTTCCTGCAACAGTGCC | | | |
| TCATGGTCATAGCTAGAACGTGGACTCCGCAGAAGATAAAAC | | | |
| TCGACTCTAGAGGAAGGGCGATGGCCCAAGCCCTAAAACATC | | | |
| GTTGTAAAACGACGTTTTGGGGTCGAGGAATATTTTTGAATG | | | |
| GGCGATTAAGTTGGAAGGGAGCCCCCGAGAACCCTTCTGACC | | | |
| CTTCGCTATTACGCACGTGGCGAGAAAGACACGACCAGTAAT | | | |
| GAGAAGTGTTTTTAGTCGGATTCTCCGTAAATGTGAGCGAGT | | | yes (edge) |
| ATTATTTACATTGGAATTAATTACATTTCGTTATACAAATTC | | | |
| CATGGAAATACCTAATGGAAACAGTACACGGAATCATAATTA | | | |
| CAGAACAATATTACTGCTTCTGTAAATCACCGACCGTGTGAT | | | |
| CTGAGTAGAAGAACTCCTTGAAAACATATAGTTAATTTCATC | | | |
| CGTTGTAGCAATACGAGTCAATAGTGAATCGCAAGACAAAGA | | | |
| GAGGCCACCGAGTAACCTTTTTAACCTCGTTGGGTTATATAA | | | yes (edge) |
| ACCACCCTCATTTTCAAAGACAAAAGGGAACAAAGTTACCAG | | | |
| ACCCTCAGAACCGCGTAAATATTGACGGATCTTACCGAAGCC | | | |
| CGGAATAGGTGTATCGTCACCGACTTGAAGCCCAATAATAAG | | | |
| AGGCGGATAAGTGCCACCAGTAGCACCATGAGCGCTAATATC | | | |
| CTCAAGAGAAGGATCAATGAAACCATCGGGGAGAATTAACTG | | | |
| CCTATTATTCTGAAAATCAAGTTTGCCTTTTACAGAGAGAAT | | | |
| AGTGCCCGTATAAACGGCATTTTCGGTCGAAACGATTTTTTG | | | |
| GTACTGGTAATAAGTTTCATAATCAAAATTACAAAATAAACA | | | |
| GTTCCAGTAAGCGTCGCCTCCCTCAGAGTACCAACGCTAACG | | | |
| GCCTGTAGCATTCCCAACATATAAAAGAGCAGTATGTTAGCA | | | |
| CTCATTAAAGCCAGAGCCACCACCCTCATAGTTGCTATTTTG | | | |
| CAGGTCAGACGATTCGCCGCCAGCATTGACCTCCCGACTTGC | | | |
| GTCAATAGATAATACAACTCGTATTAAAAAGGCTTATCCGGT | | | |
| TAAAATATCTTTAGAAGTTTGAGTAACAAGGAATCATTACCG | | | |
| AGTTGGCAAATCAACAGAAGGAGCGGAACGCACTCATCGAGA | | | |
| TTGCTGAACCTCAAATGGCAATTCATCAGTCTTTCCTTATCA | | | |
| ACGCTGAGAGCCAGTCTGAATAATGGAAATCCTAATTTACGA | | | |
| AGAGGTGAGGCGGTTTGCACGTAAAACATATCAACAATAGAT | | | |
| GCCATTAAAAATACGTTTAACGTCAGATGACAATAAACAACA | | | |
| GCTATTAGTCTTTACGGGAGAAACAATAATAAAGTACCGACA | | | |
| TGTACCGTAACACTTTTTGTCACAATCAGGAATACCCAAAAG | | | |
| TGAAAGCGTAAGAAAAGTTACAAAATCGGTAATTTAGGCAGA | | | |

| |
|---|
| AAAAGGGACATTCTCCTGAGCAAAAGAATAGGGCTTAATTGA |
| TTACCAGTATAAAGCGGTAATCGTAAAAATCGGTGCGGGCCT |
| CTAGAAAAAGCCTGTGATAATCAGAAAAGCGCCATTCGCCAT |
| AAATAAGGCGTTAAAAATATTTAAATTGCCAGCTTTCCGGCA |
| TTCTGACCTAAATTATTAAATTTTTGTTGGGGACGACGACAG |
| ACGCGAGAAAACTTACGCCATCAAAAATGGTGTAGATGGGCG |
| AAGGAAACCGAGGACGTCATAAATATTCAACTAATGCAGATA |
| CTATATGTAAATGCCTTTCATCAACATTGGGAACAAACGGCG |
| CTTTTTAAGAAAAGGTTCAGAAAACGAGGGTAGAAAGATTCA |
| AGCAAGAAACAATGTACCCTGACTATTAATCTACGTTAATAA |
| AGAGAGATAACCCAAAAGATTAAGAGGAAAGAACTGGCTCAT |
| AACACCCTGAACAATAATTCGAGCTTCATAATTTCAACTTTA |
| AACATAAAAACAGGACAGGTCAGGATTAGAGAAACACCAGAA |
| TTTAACGTCAAAAAAAGAGGTCATTTTTCGTAACAAAGCTGC |
| GCCATATTATTTATTATAATGCTGTAGCGAGTAATCTTGACA |
| AGCGTCTTTCCAGATACGGTGTCTGGAACGGTGTACAGACCA |
| CACCCAGCTACAATAATTCTGCGAACGATAAGGGAACCGAAC |
| GGGAGGTTTTGAAGTTCGCAAATGGTCACTCCATGTTACTTA |
| ATTCTAAGAACGCGGGGCGCGAGCTGAATTGTATCATCGCCT |
| CGCCCAATAGCAAGTAGCATTACATCCACGGGGAGAGGCGGT |
| ACAAGCAAGCCGTTAGCAAAATTAAGCAGAAACCTGTCGTGC |
| TTCCAAGAACGGGTGGTTGTACCAAAAATCACATTAATTGCG |
| GCATGTAGAAACCAAGAAGCCTTTATTTGCATAAAGTGTAAA |
| AACTGGCATGATTATAGTAAAATGTTTAAGTAAGAGCAACAC |
| AAGTCCTGAACAAGCTCATATATTTTAAATTGTTATCCGCTC |
| TGTTCAGCTAATGCAGATTCAAAAGGGTGCTCGAATTCGTAA |
| AAAGGTAAAGTAATATCAATATGATATTTGCATGCCTGCAGG |
| GGCATTTTCGAGCCGGAGAGGGTAGCTATTCCCAGTCACGAC |
| GAATCGCCATATTTGTCATTGCCTGAGAGGGATGTGCTGCAA |
| CTGAGGCTTGCAGGAGGCTTGCCCTGACGAGAGTACCTTTAA |
| AAATCGGAACCCTAGTAACGCCAGGGTTTTTTTGAGAGATCT |
| ATATGTACCCCGGTTTTAGTATCATATGAACAATTTCATTTG |
| AAGATTGTATAAGCATAAGAATAAACACTAAATCAATATATG |
| TTGTTAAAATTCGCTAATGGTTTGAAATGTCGCTATTAATTA |
| TTTAACCAATAGGATTTCAAATATATTTGCGATAGCTTAGAT |
| CTTCCTGTAGCCAGTGATGCAAATCCAATTTATCAAAATCAT |
| AAATGCTTTAAACATAAGCAGATAGCCGCGACATTCAACCGA |
| AAAAATCAGGTCTTAAATAGCAATAGCTAAATTATTCATTAA |
| GCGGATTGCATCAACAAGAATTGAGTTAGCCATTTGGGAATT |
| GGAAGCAAACTCCAGAAGCGCATTAGACATAGCAGCACCGTA |
| TTGCTCCTTTTGATTGAAAATAGCAGCCTTAGCGTCAGACTG |
| GCTTAATTGCTGAACCCAATCCAAATAAATAGCCCCCTTATT |
| ATATGCAACTAAAGGCCTAATTTGCCAGTCACCGGAACCAGA |
| AACAGTTGATTCCCTTTATCCTGAATCTCCGCCACCCTCAGA |
| GCCAGAGGGGGTAAAGACTCCTTATTACAACGCAAAGACACC |
| TATATTTTCATTTGAGGCGTTTTAGCGAACAGGAGTAGACTT |
| TCTACTAATAGTAGCAAATCAGATATAGTCCTTTGCCCGAAC |
| CAAGGCAAAGAATTTTTATTTTCATCGTTTATCATTTTGCGG |
| CATAAAGCTAAATCATTAAACCAAGTACTTATCATCATATTC |
| AATACTTTTGCGGGATCAATAATCGGCTATATAATCCTGATT |
| TAATGTGTAGGTAAAGAACGCGCCTGTTGAAATAAAGAAATT |
| ACAGTCAAATCACCTCTGTCCAGACGACGAATATACAGTAAC |
| GATAAATTAATGCCAGTAATAAGAGAATACGGATTCGCCTGA |
| CAATACTGCGGAATAACGCAATAATAACATAGAAAATTCATA |
| ACAAAGGCTATCAGAACAACGCCAACATCGCAGAGGCGAATT |
| AGAGAATCGATGAACCAACGCTCAACAGGATGATGAAACAAA |
| GAAAGGAGCGGGCGTGTTGGGAAGGGCGCTAGCATGTCAATC |
| ACAGGGCGCGTACTAGATCGCACTCCAGTAAACGTTAATATT |
| CGATTAAAGGGATTGGATAGGTCACGTTAATTCGCGTCTGGC |

| | | | |
|---|---|---|---|
| TAATTTTTTCACGTACAACATTATTACAAATGACCATAAATC | | | |
| TGAATTTCTTAAACCCTTATGCGATTTTAGCCCGAAAGACTT | | | |
| CGGCTACAGAGGCTCTGACCTTCATCAATCAACATGTTTTAA | | | |
| GCACCAACCTAAAACAGACGGTCAATCAGTAGATTTAGTTTG | | | |
| AGTACAAGGTTTTTCCAGGGTCGGAGATAAGGTGGCATCAAT | | | |
| GGTCCACGCTGGTTCGCTTTCCAGTCGGATAAAGCCTCAGAG | | | |
| ATAGCCCGAGATAGCATACGAGCCGGAACAACGCAAGGATAA | | | |
| TTCTGTATGGGATTGAATTACGAGGCATGACTGGATAGCGTC | | | |
| AAAAACCGTCTATCTCCCCGGGTACCGAGAGAAAGGCCGGAG | | | |
| AGCGAGAGGCTTTTATAAAAACCAAAAT | | | |
| TAGTTAGCGTAACGACAGACAGCCCTCA | | | yes (edge) |
| ATACATAAAGGTGGAACGTAGAAAATAC | | | yes (edge) |
| CAAATATCGCGTTTAGTCAGAGGGTAATTTACCATTAGCAAG | | | |
| ACCATTAGATACATCCTTAAATCAAGATGAGCCGCCACCAGA | | | |
| AAATTTTTAGAACCAAAAATAATATCCCGGGTTAGAACCTAC | | | |
| GCTGCGCGTAACCAGGAAACCAGGCAAAGCCCCAAAAACAGG | yes | | |
| TGCTTTCCTCGTTAATCTGCCAGTTTGAAAATCAGCTCATTT | yes | | |
| GGAGTGAGAATAGAGGAATACCACATTCATTGAATCCCCCTC | yes | | |
| AGGAGCCTTTAATTCGTTGGGAAGAAAATAGTCAGAAGCAAA | yes | | |
| TGACAACAACCATCGGCTTGAGATGGTTAAGCGAACCAGACC | yes | | |
| ATCTAAAGTTTTGTTTTACCAGACGACGGCAAAAGAAGTTTT | yes | | |
| CACCCTCAGCAGCGATTACCCAAATCAAGCGGATGGCTTAGA | yes | | |
| AGTTTCCATTAAACAGAGGACAGATGAAGTTTCATTCCATAT | yes | | |
| ACTCATCTTTGACCAAATCCGCGACCTGATAACCTGTTTAGC | yes | | |
| GATTGCCCTTCACCAATCGGCCAACGCGATAAATCATACAGG | yes | | |
| TGGTGGTTCCGAAAATGAGTGAGCTAACCATTATGACCCTGT | yes | | |
| GAGTCCACTATTAAGTTTCCTGTGTGAAATGCAATGCCTGAG | yes | | |
| ACCCAAATCAAGTTGCCAGTGCCAAGCTCAACCGTTCTAGCT | yes | | |
| GGGAAAGCCGGCGACAGCTGGCGAAAGGGTCTGGAGCAAAC | yes | | |
| ACGGAATAAGTTTAGAGTTTCGTCACCATTAGTAAATGAATT | | | |
| TGGTTTACCAGCGCCAGGGATAGCAAGCTCAACAGTTTCAGC | | | |
| TTGAGGGAGGGAAGCACCCTCAGAACCGGGAATTGCGAATAA | | | |
| AGGTGAATTATCACCACCGTACTCAGGAAAAAAGGCTCCAAA | | | |
| AGAGCCAGCAAAATCGTCGAGAGGGTTGGCTTGCTTTCGAGG | | yes (center) | |
| GCCGGAAACGTCACTAGGATTAGCGGGGAGTTGCGCCGACAA | | | |
| ATCAGTAGCGACAGACATGAAAGTATTAGATATATTCGGTCG | | yes (center) | |
| TAGCGCGTTTTCATCAGTTAATGCCCCCTTTTGCGGGATCGT | | | |
| AGCGTTTGCCATCTTTTTAACGGGGTCAAACGAGGGTAGCAA | | yes (center) | |
| GCCACCACCGGAACCATACATGGCTTTTCTTTTTCATGAGGA | | | |
| ACCGCCACCCTCAGAATGGAAAGCGCAGATGCCACTACGAAG | | yes (center) | |
| ACCACCACCAGAGCGGCCTTGATATTCAGAATACACTAAAAC | | | |
| TACAAACAATTCGACATTTGAGGATTTACAAGCGCGAAACAA | | yes (center) | |
| GTTATTAATTTTAAGAGCACTAACAACTGACGGGCAACAGCT | | | |
| AACAAAGAAACCACCAGTTGAAAGGAATGAGTTGCAGCAAGC | | yes (center) | |
| CTGATTATCAGATGATATCAAACCCTCAAAAATCCTGTTTGA | | | |
| GTTTGGATTATACTCAGCAAATGAAAAATATAAATCAAAAGA | | yes (center) | |
| CATATCAAAATTATCAGTATTAACACCGCCAGTTTGGAACAA | | | |
| GCGTAGATTTTCAGCGAACGAACCACCAAACGTCAAAGGGCG | | yes (center) | |
| AGTACCTTTTACATATGCGCGAACTGATCTACGTGAACCATC | | | |
| TTGCTTTGAATACCTACGTGGCACAGACTGCCGTAAAGCACT | | | |
| ATTCATTTCAATTAGGCCAACAGAGATATTTAGAGCTTGACG | | | |
| CATCAAGAAAACAACAGATTCACCAGTCGAAGGGAAGAAAGC | | | |
| AATTACCTTTTTTACATTTTGACGCTCAAGTGTAGCGGTCAC | | | |
| TGAGTGAATAACCTCGCCAGCCATTGCACTTAATGCGCCGCT | | yes (center) | |
| ATTTTCCCTTAGAATCAAACTATCGGCCGAGCACGTATAACG | | | |
| TAAGACGCTGAGAATTCTTTGATTAGTAGCTAAACAGGAGGC | | yes (center) | |
| AGGTCTGAGAGACTAAAGAGTCTGTCCATACGCCAGAATCCT | | | |

Table S5. List of DNA oligos for DNA nanotiles

| DNA nanotile sequence | biotin sites | 10 binding sites | 4 edge binding sites |
|---|---|---|---|
| TCAGGAGGTTTAGTACCGCCACCCTCAGAAC | | | |
| TTTGCTAAGTAAATGAATTTTCTGAGTGCCTT | | | |
| GAGTAACAGTTTTAACGGGGTCTATGGGAT | | | |
| TAACCGATATGACAACAACCATCGCACCACCC | | | |
| TCAGAGCCGCCACCCTCAGAGCCCCACGCA | | | |
| CACTAAAATAAAACGAAAGAGGCAACGTCACC | | | |
| AATGAAATTAGCAAGGCCGGAAAAAGAATA | | | |
| TGACCTTCTACAGACCAGGCGCATACCACGGA | | | |
| ATAAGTTAAGAAACGCAAAGACAGGCTGGC | | | |
| AAAATCTAATACCAGTCAGGACGTGCAAGAAA | | | |
| CAATGAAAGCCCAATAATAAGATGGGAAGA | | | |
| GGGGTAATTTTGCAAAAGAAGTTTAAATAAAC | | | |
| AGCCATAAATTTGCCAGTTACATGCCAGAG | | | |
| TTCGAGCTAAGACTTCAAATATCGATCGTAGG | | | |
| AATCATTGCCGTTTTTATTTTCCGTTTTAA | | | |
| AGTAGATTAGTTGATTCCCAATTCTTCTGTCC | | | |
| AGACGACCAAAAGGTAAAGTAATGCGAACG | | | |
| CCCTGTAATCGGTTGTACCAAAAATAAGGCGT | | | |
| TAAATAACGACCGTGTGATAAACATTATGA | | | |
| AGGCTATCTAGCTATTTTTGAGAGAAGACGCT | | | |
| GAGAAGAGCGATAGCTTAGATTATCTACAA | | | |
| ACGCCATCCAGCTCATTTTTTAACAGGCGAAT | | | |
| TATTCATTACAAAATCGCGCAGCAATAGGA | | | |
| AGCTTTCCCCTCAGGAAGATCGCACATCAATA | | | |
| TAATCCTCAGATGATGGCAATTCTCCAGCC | | | |
| GGTCGACTCAGTGCCAAGCTTGCAGGAGCACT | | | |
| AACAACTTCTAAAATATCTTTATGCCTGCA | | | |
| TTAATGAAGGGAAACCTGTCGTGCATTAAAAA | | | |
| TACCGAACCCTAAAACATCGCCCAGCTGCA | | | |
| CCCGAGATATCCCTTATAAATCAAAAACGCTC | | | |
| ATGGAAACCATTGCAACAGGAAAAGAATAG | | | |
| GAGCTTGACGGGGAAAAAAGGGATTTTAGAC | | | |
| TAGGAACCCATGTACACAACGCC | | | |
| TGTAGCATAATTTTTTCACGTTCCTTTAAT | | | |
| TGTATCGAGCGAAAGACAGCATTGAGGACT | | | |
| AAAGACTCATCGCCTGATAAATTTAGCCGG | | | |
| AACGAGGATTCAGTGAATAAGGTAAATTGG | | | |
| GCTTGAGTAGGAATACCACATTTTACGAGG | | | |
| CATAGTACCCCTCAAATGCTTTCAAAAATC | | | |
| AGGTCTTTCCTTTTGATAAGAGCTGAATAT | | | |
| AATGCTGGGGCGCGAGCTGAAATTAACATC | | | |
| CAATAAATTTAAATGCAATGCCGAGAAAGG | | | |
| CCGGAGAATGTCAATCATATGTCAGGAAGA | | | |
| TTGTATAACAACCCGTCGGATTTGGGATAG | | | |
| GTCACGTTGGGAAGGGCGATCGGAAAGGGG | | | |
| GATGTGCAATTGTTATCCGCTCAAAGTGTA | | | |
| AAGCCTGAGTGAGACGGGCAACGAGTTGCA | | | |
| GCAAGCGCAAAGGGCGAAAAACCATCACCCAAATCAA | | | yes (edge) |
| GCGAATAATTCCACAGACAGCCCTGGGATAGCAAGCCCAA | | | yes (edge) |
| CCCTCAGCGTTTATCAGCTTGCTTAAGGAATT | yes | | |
| ATTTGTATTTTTCATGAGGAAGTTGATCGTCA | | | |
| AGCTGCTCCGCAGACGGTCAATCACAACGGAG | yes | | |
| TTGAGATTATGGTTTAATTTCAACCGTAACAA | | | |
| ATTGAATCAGAGCAACACTATCATTTCATCAG | yes | | |
| TTAATTGCTACCCTGACTATTATAAAATATTC | | | |
| TTCATTTGTAGCTCAACATGTTTTGAGTACCT | yes | | |
| TCATATATTCATACAGGCAAGGCAGCTATATT | | | |
| AAACTAGCCAGTCAAATCACCATCTAGAACCC | yes | | |
| AGCGAGTAAGCAAATATTTAAATTTAATCGTA | | | |
| GCAACTGTTGGTGTAGATGGGCGCTAAATGTG | yes | | |
| CTGTGTGATGCAAGGCGATTAAGTCAGGCTGC | | | |
| TTTTCACCGGGTGCCTAATGAGTGGCTGTTTC | yes | | |
| TCCAACGTGTCCACGCTGGTTTGCGGTTTTTC | | | |
| GTTTTTTGGGGTCGAGACGTGGAC | | | |
| ACCACCCTCATTTTCACATAGTTAGCGTAACGTAGAAAGG | | | |

| | | |
|---|---|---|
| AACAACTATCGAGGTGAATTTCTTAAGGCCGC | | yes (center) |
| TTTTGCGGTCCATTAAACGGGTAAAGCGCGAA | | |
| ACAAAGTATAAGGGAACCGAACTGTCATTACC | | |
| CAAATCAATTTAATCATTGTGAATATTACAGG | | |
| TAGAAAGAAACCCTCGTTTACCAGACTGCGGA | | |
| ATCGTCATGTCAGAAGCAAAGCGGACAGGTCA | | |
| GGATTAGAAAATATGCAACTAAAGGTCAATAA | | yes (center) |
| CCTGTTTAAAGAATTAGCAAAATTCAAGGATA | | |
| AAAATTTTAATATGATATTCAACCGAGAATCG | | |
| ATGAACGGGTAAACGTTAATATTTCAGCTTTC | | |
| ATCAACATATCGTAACCGTGCATCAGCGCCAT | | |
| TCGCCATTTGGGTAACGCCAGGGTCGTAATCA | | |
| TGGTCATAAGCTAACTCACATTAATATTGGGC | | yes (center) |
| GCCAGGGTCCCAGCAGGCGAAAATAGTCCACT | | |
| ATTAAAGAGTGCCGTAAAGCACTA | | |
| AGTGAGAAATCTAAAGTTTTGTCGCCACCCTCAGAGCC | | |
| GGGAGTTAAAACAGCTTGATACCTCAGCGG | | |
| TTATACCAAATACGTAATGCCACGCTTGCA | | |
| CCGGATATACCAACTTTGAAAGACCAGCGA | | |
| ACAACATTTACCTTATGCGATTTACAAGAA | | |
| CGTCCAATACGACGATAAAAACCTAACGGA | | |
| AAACTCCAATTGCATCAAAAAGATGGATAG | | |
| CGCAAATGTACGGTGTCTGGAAGCGGAAGC | | |
| TTTCAACGAAGCAATAAAGCCTCTACATTT | | |
| GCAAACAAGTTCTAGCTGATAAAGCCTTTA | | |
| CCTGTAGCTGTTAAAATTCGCATGTCTGGA | | |
| CCAGGCAATGCCAGTTTGAGGGGTGGCCTT | | |
| CTCGAATTTTTCCCAGTCACGACCCGGAAA | | |
| GGTTTGCGTTGCGTTGCGCTCACTACCGAG | | |
| GGAACAAGCCTGTTTGATGGTGGGAGAGGC | | |
| AATCGGAACCCTAAACCAGTTT | | |
| CGCCACCCTCAGAACCGTCTTTCCAGACGTTAACAACTTT | | |
| CAACAGTTGATAGTTGCGCCGACAATATTCGG | | |
| TCGCTGAGTACGAAGGCACCAACCCACTCATC | | |
| TTTGACCCGGACAGATGAACGGTGATCAAGAG | | |
| TAATCTTGTAAGAACTGGCTCATTCGTTAATA | | |
| AAACGAACAAAATAGCGAGAGGCTAGTAAAAT | | |
| GTTTAGACTTAAGAGGAAGCCCGATCAAAGCG | | yes (center) |
| AACCAGACTTTCATTCCATATAACTAGTTTGA | | |
| CCATTAGAAGAGCATAAAGCTAAATACTTTTG | | |
| CGGGAGAATTAATGCCGGAGAGGGAGGTCATT | | |
| GCCTGAGATAAATTTTTGTTAAATAAAAATAA | | yes (center) |
| TTCGCGTCACGACGACAGTATCGGGGCACCGC | | |
| TTCTGGTGGTTGTAAAACGACGGCCTAGAGGA | | |
| TCCCCGGGTGCCCGCTTTCCAGTCTCGGCCAA | | |
| CGCGCGGGTTCCGAAATCGGCAAAAGGGTTGA | | |
| GTGTTGTTGGGAGCCCCCGATTTA | | |
| TGGTAATAAGTGCCCGTATAAACAAGGTGTATCACCGTAC | | |
| CTCAGAACCGCCACCAGAACCACCGAGTGTAC | | |
| CATTACCACCATCGATAGCAGCACCCGCCACC | yes | |
| ACATATAATATTTTGTCACAATCAAGTAGCAC | | |
| TTGAGTTAATAGCAATAGCTATCTAGGTGGCA | yes | |
| CAGAGCCTTTATTTATCCCAATCCCACAAGAA | | |
| ACAAGCAAACCGCGCCCAATAGCACGTCTTTC | yes | |
| AGTACCGAGACAATAAACAACATGCATCGAGA | | |
| TGAAATACGAATAAACACCGGAATGAATATAA | yes | |
| AAAACATAGTCAATAGTGAATTTATAATGGTT | | |
| TACCAAGTTTCAATTACCTGAGCAAATCCTTG | yes | |
| CTGATTATGATTGTTTGGATTATAGCTTTGAA | | |
| GAAGGTTAAATAGATTAGAGCCGTTCATATTC | yes | |
| ACTGATAGCGAACCACCAGCAGAAGAATTGAG | | |
| ACCGCCAGTACCTACATTTTGACGATGCGCGA | yes | |
| AGGAACGGTACGCCAGACAATATT | | |
| AAGTATAGCCCGGAATGTTAATGCCCCCTGCCGCTTTTGA | | |
| TGATACAGACCAGAGCCGCCGCCAACCGCCTC | | |
| CCTCAGAGCGTAATCAGTAGCGACAGCCAGCA | | |
| AAATCACCATAGAAAATTCATATGGAAAATAC | | |
| ATACATAATACCGAAGCCCTTTTTTATCAGAG | | |
| AGATAACCAAATAAGAAACGATTTTACCAACG | | |

| | | | |
|---|---|---|---|
| CTAACGAGAGCAAATCAGATATAGAACCAAGT | | yes (center) | |
| ACCGCACTTTCAGCTAATGCAGAACGAGCCAG | | | |
| TAATAAGACATAATTACTAGAAAATCTTCTGA | | | |
| CCTAAATTTCAAAATCATAGGTCTATTAATTT | | | |
| TCCCTTAGAAAGAAGATGATGAAAACGGATTC | | yes (center) | |
| GCCTGATTCTTCTGAATAATGGAAAGGAGCGG | | | |
| AATTATCACAATAGATAATACATTAATCAACA | | | |
| GTTGAAAGGATAAAACAGAGGTGATGGCTATT | | | |
| AGTCTTTACTCAATCGTCTGAAATGCTGGTAA | | | |
| TATCCAGAAATCCTGAGAAGTGTT | | | |
| CATACATGTATTTCGGAACCTATCGAGAGGGTTGATAT | | | |
| CCACCGGAGCATTGACAGGAGGTTAAGCGT | | | |
| GGAATTAGAGAATCAAGTTTGCCAGAGCCA | | | |
| CAAACGTAGTTTACCAGCGCCAACCATTTG | | | |
| AGCGCTAAAAGAAAAGTAAGCAGATGTTAG | | | |
| CTGAATCTTTTGTTTAACGTCAAGTAATTG | | | |
| GGGTATTAAAGGCTTATCCGGTATTTTATC | | | |
| GGCATTTTCGCGCCTGTTTATCACAAGAAC | | | |
| TAATTTCAAGCCTGTTTAGTATCAGGCAGA | | | |
| CGCTATTAGAGAGACTACCTTTTTTTTAGT | | | |
| AAACAATACAAACATCAAGAAAAAAAATCGT | | | |
| CCACCAGAGGGTTAGAACCTACCTCGGGAG | | | |
| AGTTGGCATGAGGATTTAGAAGTAAAGAAA | | | |
| TTTTTGAAGGCGGTCAGTATTAATCTGGTC | | | |
| TCGGCCTTGGATTATTTACATTGGACAATA | | | |
| TTTATAATCAGTGAGCAAACTA | | | |
| GGCGGATAAGTGCCGTTATTCTGAAACATGAAGAATTTAC | | | |
| CGTTCCAGTGAGGCAGGTCAGACGCAAAATCA | | yes (center) | |
| CCGGAACCTTTAGCGTCAGACTGTCCGTCACC | | | |
| GACTTGAGAGACAAAAGGGCGACACTCCTTAT | | | |
| TACGCAGTATAGCCGAACAAAGTTTGAACAAA | | | |
| GTCAGAGGAAATGAAAATAGCAGCTTGCACCC | | | |
| AGCTACAATTCTAAGAACGCGAGGTCTTTCCT | | | |
| TATCATTCACAATAGATAAGTCCTCGCCAACA | | yes (center) | |
| TGTAATTTATATGCGTTATACAAAAACTTTTT | | | |
| CAAATATATAACCTCCGGCTTAGGATAACCTT | | | |
| GCTTCTGTCAAAATTAATTACATTAACAGTAC | | | |
| CTTTTACAATATCAAAATTATTTGTATCATTT | | | |
| TGCGGAACATTAGACTTTACAAACAAACCCTC | | | |
| AATCAATACACCGCCTGCAACAGTAAGAATAC | | yes (center) | |
| GTGGCACAGCAGATTCACCAGTCACCTGAGTA | | | |
| GAAGAACTGCCACCGAGTAAAAGA | | | yes (edge) |
| CAGTCTCTAGTATTAAGAGGCTGAGTTTTGCTCAGTACCA | | | yes (edge) |
| TTCATAATATTGGCCTTGATATTCGGAAAGCG | | | |
| AATTATCAAGCGCGTTTTCATCGGGCCATCTT | | | |
| GATTAAGATTCAACCGATTGAGGGTAAAGGTG | | | |
| GAACACCCACCAGAAGGAAACCGAACTGGCAT | | | |
| TTGCTATTCTTTACAGAGAGAATAAATTAACT | | | |
| ATCGGCTGCGTTTTAGCGAACCTCAAGATTAG | | | |
| TTTAACAAGAACAAGAAAAATAATAATCAATA | | | |
| CGCGAGAATTCTTACCAGTATAAATCGCCATA | | | |
| GTGAGTGATTGGGTTATATAACTAACAAAGAA | | | |
| TATACAGTTAACAATTTCATTTGACAATATAT | | | |
| AGTAACATCACGTAAAACAGAAATCAGATGAA | | | |
| CAAATATCAATTCGACAACTCGTAAAAGTTTG | | | |
| GAAAGCGTGCCACGCTGAGAGCCACTGAACCT | | | |
| ATCACTTGCACGACCAGTAATAAATCTGACCT | | | |
| GTCTGTCCATCACGCAGTAATAAC | | | |
